# Endogenous ZAP affects Zika virus RNA interactome

**DOI:** 10.1101/2024.05.23.595534

**Authors:** Ahmad Jawad Sabir, Nguyen Phuong Khanh Le, Prince Pal Singh, Uladzimir Karniychuk

**Affiliations:** Department of Microbiology and Immunology, University of Illinois, College of Medicine, Chicago, IL, USA; Department of Veterinary Biosciences, College of Veterinary Medicine, The Ohio State University, Columbus, OH, USA; School of Public Health, University of Saskatchewan, Saskatoon, Canada

**Keywords:** Flavivirus, Zika virus, Japanese encephalitis virus, West Nile virus, Zinc finger antiviral protein, ZAP, RNA, RNA helicase, interactome

## Abstract

One of the most recent advances in the analysis of viral RNA–cellular protein interactions is the Comprehensive Identification of RNA-binding Proteins by Mass Spectrometry (ChIRP-MS). Here, we used ChIRP-MS in mock-infected and Zika-infected wild-type cells and cells knockout for the zinc finger CCCH-type antiviral protein 1 (ZAP). We characterized “ZAP-independent” and “ZAP-dependent” cellular protein interactomes associated with flavivirus RNA and found that ZAP affects cellular proteins associated with Zika virus RNA. The ZAP-dependent interactome identified with ChIRP-MS provides potential ZAP co-factors for antiviral activity against Zika virus and possibly other viruses. Identifying the full spectrum of ZAP co-factors and mechanisms of how they act will be critical to understanding the ZAP antiviral system and may contribute to the development of antivirals.

## INTRODUCTION

Emerging flaviviruses constantly threaten public health. The National Institute of Allergy and Infectious Diseases classifies flaviviruses—Zika virus, Japanese encephalitis virus (JEV), yellow fever virus (YFV), and West Nile virus (WNV) as a “Category B Priority,” the 2^nd^ highest priority of emerging pathogens and biodefense threats. Zika virus caused the human epidemic in 2015 and persists in at least 89 countries. Japanese encephalitis virus causes encephalitis in humans in the Asia-Pacific region [1], with 68,000 annual cases and 15,000 deaths [2]. There is a concern that JEV can be introduced into North America given the large population of amplifying hosts—pigs and wild boars, and susceptible *Culex* mosquitoes [3–5]. Japanese encephalitis virus keeps expanding—the 2022 JEV outbreak in Australia with infections in swine herds, zoonotic transmission, and human deaths was caused by genotype IV which was not associated with outbreaks before [6]. West Nile virus is the most common arthropod-borne virus in the US. In 2022, the US Centers for Disease Control and Prevention reported 1,126 cases and 93 deaths. In 2023, there were 2,406 human cases in the US, including 1,599 neuroinvasive diseases. The rising number of human cases caused by mosquito-borne flaviviruses shows the lack or inefficiency of environmental controls. There are no approved human vaccines for some flaviviruses (i.e., Zika, WNV). Also, there are no licensed antivirals against flaviviruses because of a knowledge gap in cellular pathways controlling infections, including the knowledge gap in interactions of viral RNA with cellular proteins.

High-throughput methods enable global analysis of viral RNA–cellular protein interactions. One of the most recent advances in analysis of viral RNA–cellular protein interactions is the Comprehensive Identification of RNA-binding Proteins by Mass Spectrometry (ChIRP-MS) [7], where interactions between cellular proteins and genomic RNA of several viruses were profiled that advanced the fundamental knowledge of RNA viruses and informed novel antivirals [8–11]. To our knowledge, all these ChIRP-MS studies were done in wild-type cell lines. And comparative ChIRP-MS studies on viral RNA–cellular protein interactions in wild-type and knockout (KO) cell lines are not reported.

The zinc finger CCCH-type antiviral protein 1, also known as ZAP, ZC3HAV1, or PARP13, is a cellular protein with broad antiviral activity. The protein was first characterized to inhibit murine leukemia virus [12] and later alphaviruses, filoviruses, influenza virus, porcine reproductive and respiratory syndrome virus, hepatitis B virus, human cytomegalovirus, human T cell leukaemia virus type 1, and human immunodeficiency virus-1 [13–23]. The previous study in A549 cells showed JEV sensitivity to ectopic and endogenous ZAP [24]. In the same study, Zika virus was not sensitive to ectopic ZAP; the sensitivity of Zika virus to endogenous ZAP was not tested [24]. Another study using ZAP wild-type and ZAP knockout A549 cells showed that wild-type Zika virus was also not sensitive to antiviral ZAP effects [25]. However, our recent study revealed that Zika virus was sensitive to endogenous ZAP in VERO cells. Specifically, we observed reduced viral RNA, infectious titers, and Zika virus NS5 protein expression in wild-type cells compared to ZAP knockout cells [26]. This discrepancy may be due to the different cell lines used: previous studies utilized A549 cells derived from human lung cancer tissue [24], while we used VERO cells from a healthy monkey. Antiviral signaling for many cellular proteins can be highly cell-specific [27]. Another factor is the species difference: the previous study used a human cell line, and we used an African green monkey cell line, a possible natural reservoir host of Zika virus [28]. Species-specific ZAP effects are documented [29,30], but comparative studies on ZAP antiviral activity between primates are lacking.

ZAP binds viral RNA and evokes antiviral activity by mediating viral RNA degradation and translational inhibition [31]. It also interacts with various cellular proteins that may act as co-factors, enhancing its antiviral effects [32]. Furthermore, ZAP can augment other antiviral systems [14,26]. To better understand ZAP–viral RNA–cellular protein interactions, here we used Zika virus infection in wild-type and ZAP knockout VERO cells as a model. Specifically, we used ChIRP-MS in mock-infected and Zika-infected wild-type and ZAP knockout cells to characterize “ZAP-independent” and “ZAP-dependent” cellular protein interactomes associated with flavivirus RNA. Our findings suggest that ZAP influences cellular proteins associated with Zika virus RNA.

## RESULTS

### Endogenous ZAP affects flavivirus infection phenotypes in VERO cells

While Zika virus was designated as ZAP-resistant in A549 cells [24,25], in our recent study, we showed that Zika virus is sensitive to endogenous ZAP in VERO cells with reduced levels of viral RNA, infectious titers, and NS5 protein expression [26]. Previously we used the multiplicity of infection (MOI) of 1 for comparative studies in VERO wild-type (VERO-ZAP-WT) and knock-out (VERO-ZAP-KO) cell lines [26]. Here to further confirm antiviral ZAP effects against Zika virus we conducted comparative studies in the same cells with low MOI 0.01. We also included into the comparative study JEV previously shown to be sensitive to endogenous and ectopically overexpressed ZAP, and WNV with unknown sensitivity to ZAP in VERO cells.

For all flaviviruses, we observed a significant (*p* < 0.0001) reduction in E protein expression in VERO-ZAP-WT compared to VERO-ZAP-KO cells (**Figs 1A**, **B**, **E**, **F**, **I**, **J**, and **Figs S1, S2, S3**). All three flaviviruses also showed a considerable reduction of NS5 protein expression in VERO-ZAP-WT compared to VERO-ZAP-KO cells (**Figs 1D**, **H**, **L**). West Nile virus showed comparable intracellular viral RNA loads (p = 0.1875) in both VERO-ZAP-WT and VERO-ZAP-KO cells (**Fig 1K**). Consistent with the reduction of E and NS5 protein expression, Zika virus and JEV had significantly higher intracellular RNA loads in VERO-ZAP-KO cells than in VERO-ZAP-WT cells (p = 0.0012 and 0.0007; **Figs 1C, G**).

**Fig 1.**
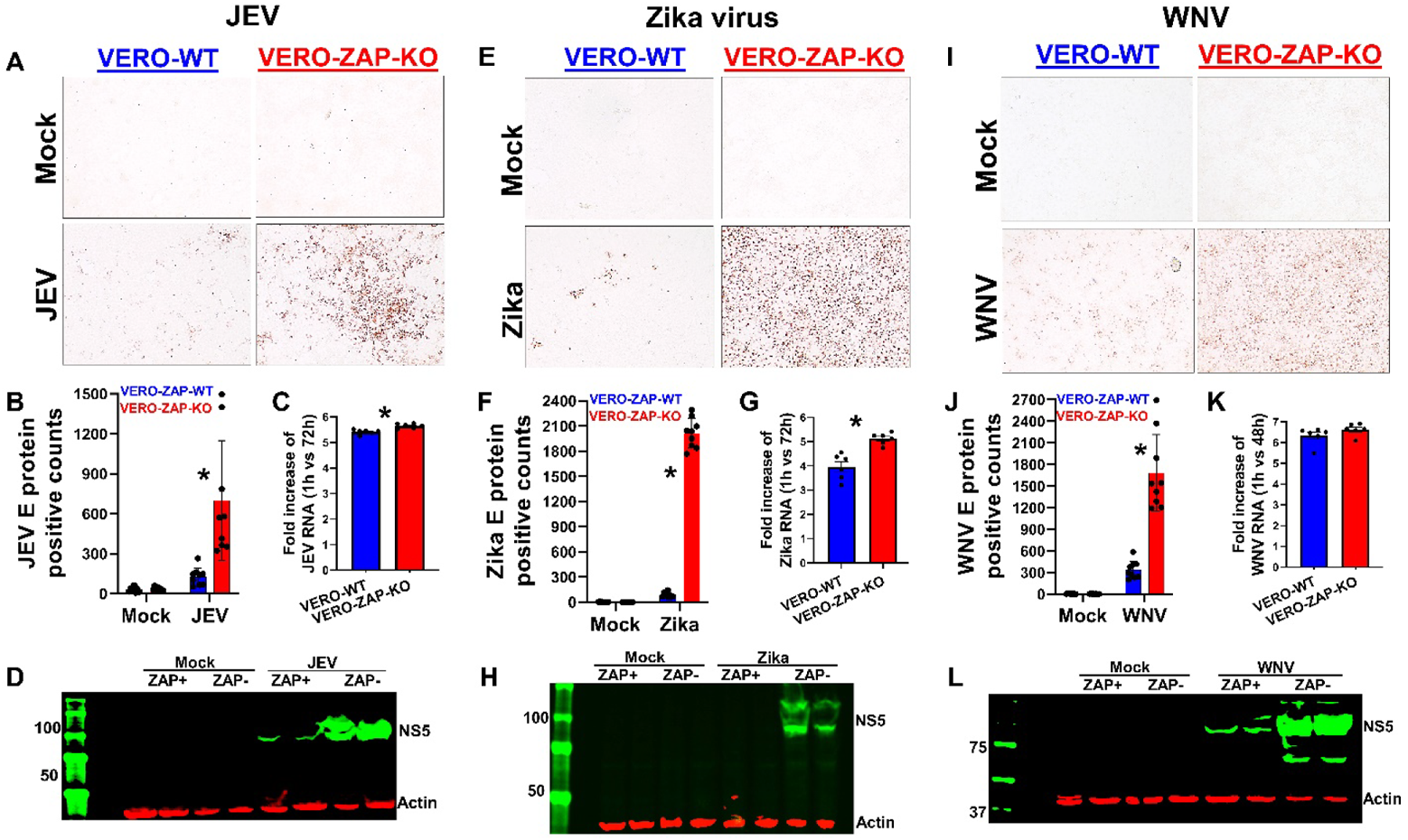
ZAP affects JEV, Zika, and WNV infection phenotypes. Representative images of cells positive for JEV (**A**), Zika virus (**E**), and WNV (**I**) E protein (red staining) at 72 h (48 h for WNV), MOI 0.01. Magnification x200. The experiment was done in 3 biological and 3 technical replicates. Figures represent the general patterns in all replicates. All 9 replicates for VERO-ZAP-WT and VERO-ZAP-KO cells are shown in **Fig S1**. The digital quantification of JEV (**B**), Zika (**F**), and WNV (**J**) positive cells in all 9 replicates from **Fig S1**. **\***Unpaired t-test: *P* < 0.05. In mock-infected cells digital sensitive counting represents the staining background. The fold increase of JEV (**C**), Zika (**G**), and WNV (**K**) RNA loads in cell lysates collected at 1 h and 72 h (48 h for WNV) after inoculation. VERO-WT and VERO-ZAP-KO cells were inoculated with MOI 0.01. Virus inoculums were removed and replaced with media. Cell lysates were collected with a lysis buffer at 1 h post-inoculation (to normalize for leftover virus inoculum RNA) and at 48-72 h. **\***Unpaired t-test: *P* < 0.05. The experiment was done in 3 biological and 2 technical replicates. (**D**, **H**, **L**) Reduced flavivirus infection in VERO-ZAP-WT cells at 72 h (48 h for WNV) represented by western blot for NS5 proteins. Normalized 50 µg of protein was used for all samples. Two biological replicates for each condition are shown.

Altogether, we confirmed ZAP antiviral effects against JEV and Zika virus, and demonstrated, for the first time to our knowledge, the sensitivity of WNV to ZAP in VERO cells. Different infection phenotypes in VERO-ZAP-KO than in VERO-ZAP-WT cells suggested ZAP effects mediated via viral RNA binding activity which may involve interactions with other cellular proteins. This motivated us to conduct a ChIRP-MS study to characterize “ZAP-independent” and “ZAP-dependent” cellular protein interactomes associated with Zika virus RNA.

### ChIRP-MS uncovers ZAP-dependent Zika virus RNA interactome

To define the ZAP-depended host protein interactome associated with Zika genomic RNA, we used the RNA-directed proteomic discovery method ChIRP-MS [7–11] in VERO-ZAP-WT and VERO-ZAP-KO cell lines. We crosslinked mock-infected and virus-infected cell lines with formaldehyde to preserve viral RNA–protein complexes and applied biotinylated oligonucleotides (**Table S1A**) tiling the entire Zika virus RNA to enrich viral RNA–host protein complexes (**Fig 2A**).

**Fig 2.**
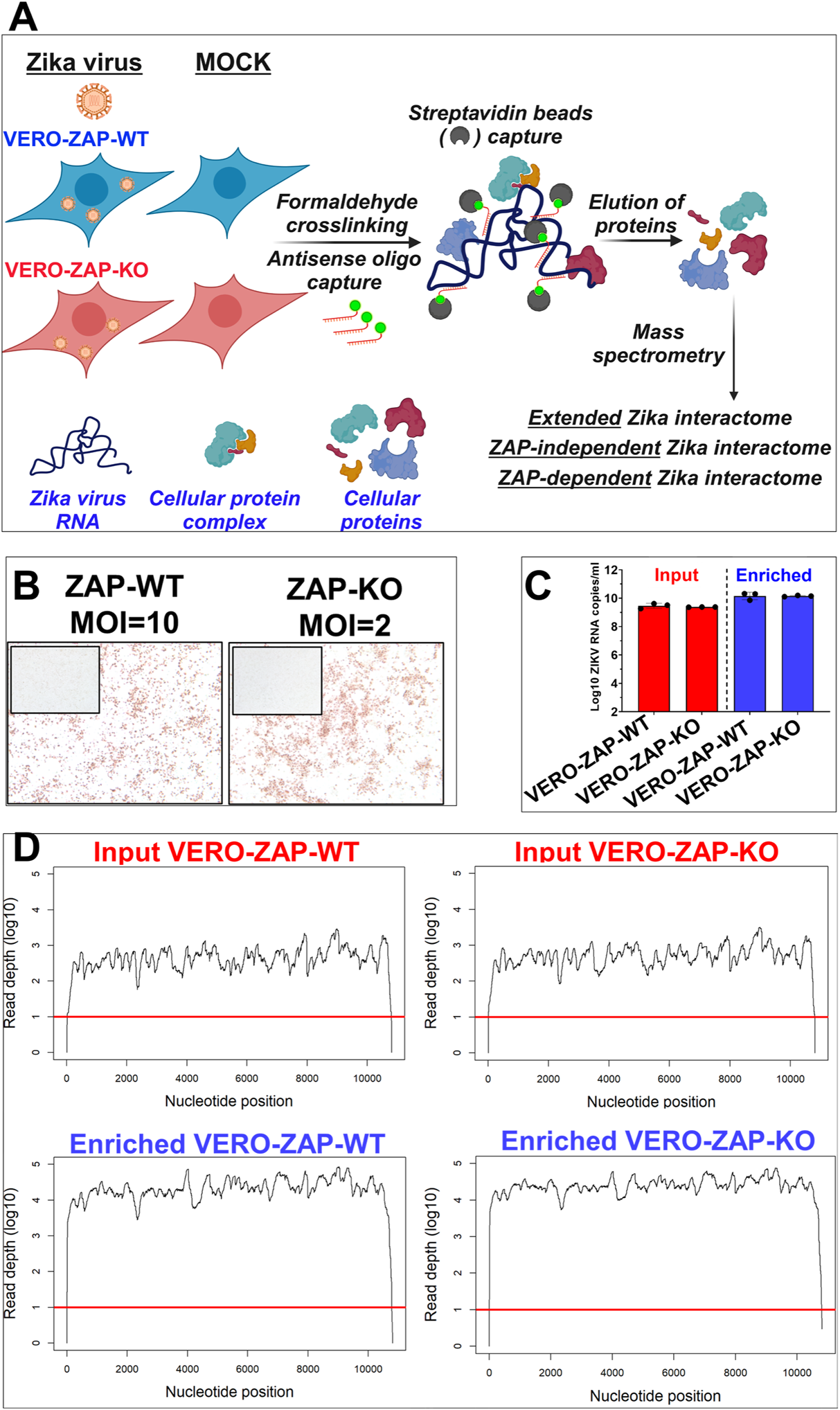
ChIRP-MS study design and validation. (**A**) ChIRP-MS experimental design to compare viral RNA–host protein interactions in ZAP-positive and ZAP-KO cells. ChIRP-MS was done in three biological replicates; each biological replicate consisted of lysed cells from four T-175 flasks for each experimental condition. **(B)** We adapted normalized MOIs to induce similar Zika viral loads in ZAP-WT and ZAP-KO cells. Red staining represents a similar number of Zika virus E protein-positive cells in WT and KO cells. Inserts—mock-infected cells. **(C)** The same Zika RNA loads in ZAP-WT and ZAP-KO cells were also confirmed using RT-qPCR. **Input** – sonicated cellular lysates. **Enriched** – immunoprecipitated viral RNA-protein complexes bound to MyOne C1 magnetic beads at the last washing step before protein elution. RNA was extracted from a 40 ul of cell lysate or bead suspension in the washing buffer representing each biological replicate and subjected to Zika virus-specific RT-qPCR; dots represent ChIRP-MS biological replicates. **(D)** The same nearly entire Zika RNA genome coverage and NGS depth in ZAP-WT and ZAP-KO cells. Representative data from one ChIRP-MS biological replicate for ZAP-WT and ZAP-KO cells. **Input** and **Enriched** are the same sample types as for RT-qPCR in **C**. Control NGS in mock-infected cells did not show Zika-specific sequences (**Fig S4**). A red line shows the 10-nucleotide NGS depth threshold.

For the Zika virus comparative study in VERO-ZAP-WT and VERO-ZAP-KO cells, we used equal MOIs for inoculation to identify ZAP effects on infection phenotypes (**Fig 1**). However, the different infection phenotypes in VERO-ZAP-WT and VERO-ZAP-KO cells (**Fig 1**) pose a challenge for ChIRP-MS study where it is essential to induce equal Zika virus RNA loads in both cell lines for an accurate comparison of ZAP-dependent and ZAP-independent interactomes. Thus, before comparing viral RNA–host protein interactions, we tested different Zika MOIs for inoculation of ZAP-WT and ZAP-KO cells to ensure similar viral RNA loads at the time of formaldehyde fixation and accurate comparative ChIRP-MS. For this, we incrementally increased the MOI for VERO-ZAP-WT cells, as Zika virus caused a more aggressive infection in VERO-ZAP-KO cells (**Fig 1**). The multiplicity of infection 10 for VERO-ZAP-WT cells and MOI 2 for VERO-ZAP-KO cells evoked similar Zika virus loads. To exclude the effects of differential uptake of the stimulus during MOI 10 inoculation of VERO-ZAP-WT cells, during VERO-ZAP-KO MOI 2 inoculation we added heat-inactivated Zika virus equivalent of MOI 8. After extensive washing and 72 h incubation, the same Zika antigen loads in VERO-ZAP-WT and VERO-ZAP-KO cells were confirmed with viral E protein staining (**Fig 2B**). The same Zika RNA loads in VERO-ZAP-WT and VERO-ZAP-KO cells were also confirmed using Zika-specific RT-qPCR and next-generation-sequencing (NGS) in sonicated cellular lysates (**input**) and immunoprecipitated viral RNA-protein complexes (**enriched**) (**Figs 2C, D; Fig S4; Supplemental Dataset 1**). High RT-qPCR Zika loads and nearly entire Zika genomic coverage with high NGS depth (**Figs 2C, D; Fig S4**) confirms that the ChIRP method efficiently recovered the same high loads of viral RNA in both VERO-ZAP-WT and VERO-ZAP-KO cells. Also, as expected, viral RNA loads were consistently and considerably (around 10^1^ difference) higher in enriched than in input samples in RT-qPCR and NGS assays (**Figs 2C, D; Fig S4**). Altogether, we confirmed normalized viral loads and comparable enrichment of viral RNA during ChIRP in VERO-ZAP-WT and VERO-ZAP-KO cells, permitting quantitative comparison of proteins associated with viral RNA between wild-type and ZAP-KO cells.

Next, we used the Mass Spectrometry Interaction STatistics (MiST) to analyze raw ChIRP-MS data (**Supplemental Dataset 2**) previously applied in ChIRP-MS studies [11,33,34]. We employed MiST with default parameters and used the Singleton Filtering option to exclude proteins with spectral counts in only one biological replicate, quantifying proteins with a MiST score of 0.75 or above [11,33,34]. Additionally, we excluded proteins that had at least one spectral count in any biological replicate of mock-infected wild-type or ZAP-KO cells [11,32]. We applied three analytical strategies to analyze ChIRP-MS data (**Table S1**):

For the first analytical strategy, using the raw ChIRP-MS data with a total of 2,269 proteins (**Table S1B**), we identified proteins specifically associated with Zika virus RNA in VERO-ZAP-WT cells. We defined the resultant list of enriched proteins as the “extended” Zika virus RNA interactome in VERO-ZAP-WT cells. This extended interactome contains 383 proteins (**Table S1C**). We named this interactome “extended” because additional control criteria were applied to narrow down and enrich the “ZAP-dependent” Zika virus RNA interactome (see below).

ChIRP-MS is a rather new technique in molecular biology; the first time it was described for systemic identification of cellular proteins interacting with cellular mRNA in 2015 [7]. From 2019, several virology groups have applied ChIRP-MS to characterize interactions between cellular proteins and genomic RNA of severe acute respiratory syndrome coronavirus 2, Ebola virus, dengue virus, and Zika virus in wild-type ZAP-positive cells [8–11]. Thus, we compared our extended Zika virus RNA interactome in VERO-ZAP-WT cells with the Zika interactome in Huh cells from a previous publication [10]. A similar number of proteins was reported here—383, and previously—395. The comparison also showed a good reproducibility of 25.4 % between two protein sets (**Table S1D**). We consider 25.4% a good reproducibility because studies were done in cells of different species—nonhuman primates (VERO) and humans (Huh), with different MOIs and sampling time points.

The specificity of our ChIRP-MS is also confirmed by physical associations between Zika virus RNA and viral prM, C, E, NS1, NS2A, NS2B, NS3, and NS5 proteins in only infected cells, while all mock-infected replicates were negative (**Table S1E**). The enrichment of viral proteins provides evidence that ChIRP-MS covers interactions across the entire length of viral RNA [8]. Analysis of the MiST abundance score and absolute spectral counts showed that the viral NS3 and NS5 proteins were most strongly recovered (**Table S1E**; NS3 MiST abundance score 0.167, NS5 MiST abundance score 0.304; NS3 mean spectral counts 39, NS5 mean spectral counts 102), which is consistent with previous ChIRP-MS Zika virus study where NS3 and NS5 were also most abundant [10]; and with direct binding function of flavivirus NS3—RNA helicase, and NS5—RNA-dependent RNA polymerase.

Analyzes of top interacting cellular proteins with the highest spectral counts and GO enrichment analysis also suggested the specificity of ChIRP-MS. All top 15 interacting proteins (except NCOA5) with the highest spectral counts (**Fig 3A**) are known factors of RNA biology, including RNA binding, RNA splicing, and RNA helicase activities (**Table S1C**). Gene Ontology enrichment analysis also showed that all seven enriched pathways with at least two-fold enrichment represent nucleic acid/RNA binding and RNA biology processes (**Fig 3D**).

**Fig 3.**
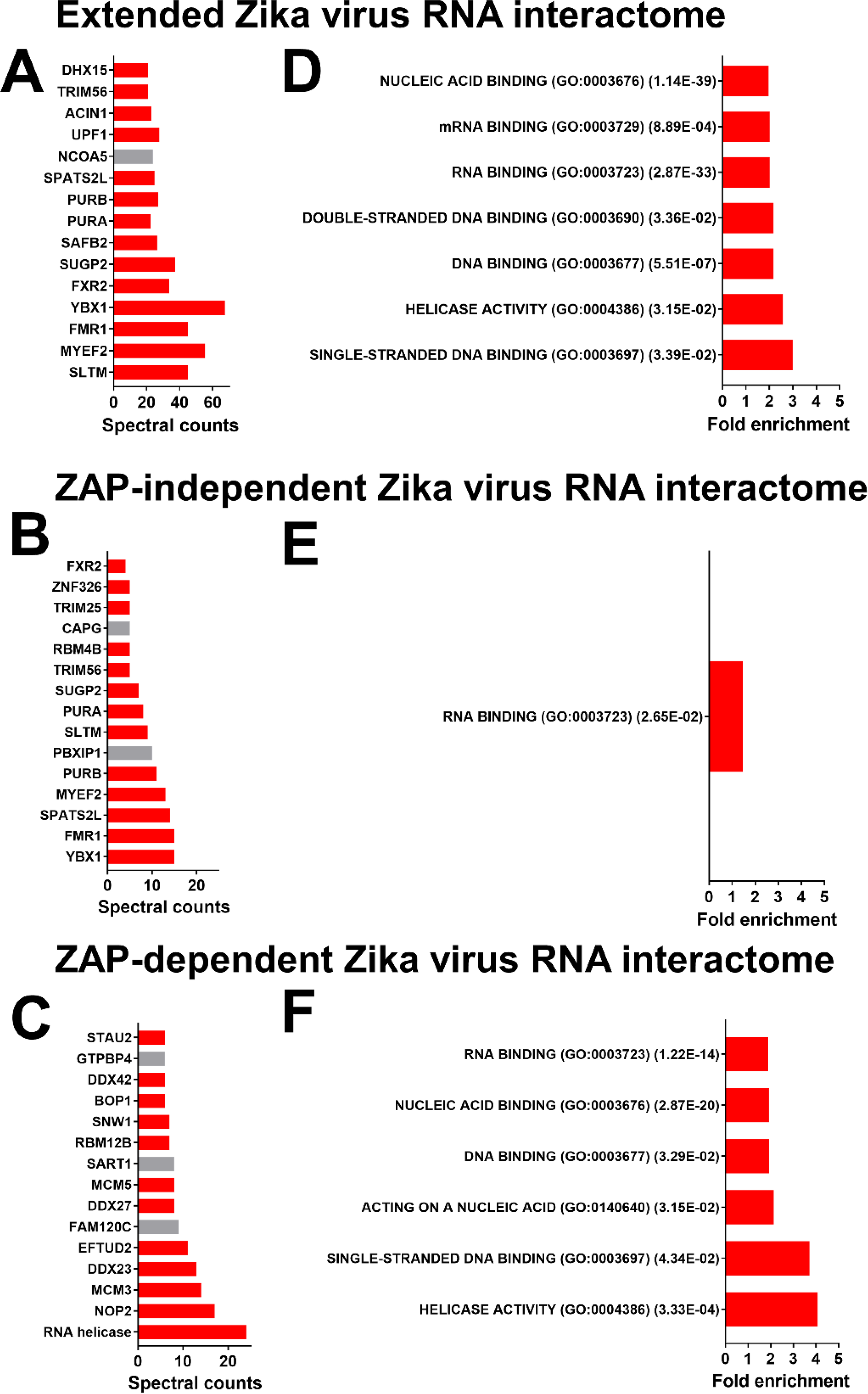
Top ChIRP-MS proteins and GO pathways identified in the Extended, ZAP-independent, and ZAP-dependent Zika virus RNA interactomes. Mean spectral counts are shown for proteins in (**A**, **B**, **C**). Spectral counts for all three biological replicates are shown in **Tables S1C, F, H**. The top proteins with unknown functions in RNA biology are highlighted in grey. For GO pathways in (**D, E, F**), GO identification numbers and FDR-adjusted *P* values are shown in parentheses. All GO processes were overrepresented.

For the second analytical strategy, we identified proteins specifically interacting with Zika virus RNA in VERO-ZAP-KO cells. For specificity, we applied the same strategy as above: We selected the Singleton Filtering MiST option to exclude proteins with spectral counts in only one biological replicate and quantify proteins with the MiST score of 0.75 or above [11,33,34]. We also excluded all proteins with a single spectral count in at least one of three biological replicates in mock-infected wild-type or ZAP-KO cells. We defined the resultant list of enriched proteins as a “ZAP-independent” Zika virus RNA interactome. We defined this interactome as ZAP-independent because associated cellular proteins were discovered in VERO-ZAP-KO cells showing the ZAP-independent nature of interactions. The ZAP-independent interactome contains 116 proteins (**Table S1F**). As expected, most, 13 out of 15 top cellular interacting proteins with the highest spectral counts are known factors in RNA biology (**Fig 3B**; **Table S1F**). Zika virus proteins were identified in only infected VERO-ZAP-KO cells (**Table S1G**).

Finally, to discover the “ZAP-dependent” Zika RNA interactome in VERO-ZAP-WT cells, in addition to the above specificity exclusion criteria in MiST settings and mock-infected cells, we excluded proteins that had at least one spectral count in at least one of the three biological replicates in VERO-ZAP-KO cells infected with Zika virus. The ZAP-dependent interactome contains 209 proteins (**Table S1H**). The ZAP-dependent Zika RNA interactome has more proteins than ZAP-independent—209 versus 116 proteins. The difference in 93 proteins between interactomes in ZAP-WT and ZAP-KO cells is rational because in the previous study a tandem mass spectrometry showed that 114 cellular proteins interact with the overexpressed immunoprecipitated ZAP large isoform in uninfected cells [32]. Other large-scale interactome studies in uninfected cells have identified more than 250 potential cellular proteins that may potentially interact with ZAP [32,35–38].

As expected, most, 12 out of 15 top interacting proteins with the highest spectral counts in the ZAP-dependent interactome are known factors in RNA biology (**Fig 3C**; **Table S1H**). Comparative GO enrichment analysis between ZAP-independent and ZAP-dependent interactomes showed only one enriched pathway related to RNA biology in ZAP-KO cells and six enriched RNA biology pathways in ZAP-WT cells, including “helicase activity” with the highest fold enrichment (**Figs 3E, F**). Accordingly, among the top 15 proteins in the ZAP-dependent interactome four proteins were RNA helicases (**Table S1H**). In total, 11 RNA helicases were enriched in the ZAP-dependent interactome—DDX42, DDX56, DDX27, DDX50, DDX52, DDX23, DDX54, DDX18, DHX57, DDX47, and one uncharacterized RNA helicase (**Tables S1H, I**).

Altogether, using ChIRP-MS and MiST analysis, we characterized the cellular interactome associated with Zika virus RNA, including Extended, ZAP-independent, and ZAP-dependent interactomes.

### ZAP wild-type and knockout VERO cells express DDX RNA helicases

ZAP binds cellular mRNA and affects gene expression, including antiviral and immune resolution transcriptional responses [39,40]. Thus, it was unclear whether the enrichment of RNA helicases in the ChIRP ZAP-dependent Zika interactome (**Tables S1H; Figs 3C, F**) and the depletion of RNA helicases in the ChIRP ZAP-independent Zika interactome (**Tables S1F; Figs 3B, E**) were exclusively due to direct ZAP–viral RNA–cellular RNA helicase interactions in ZAP-WT cells and the lack of such interactions in ZAP-KO cells, or ZAP-mediated effects on cellular RNA helicase mRNAs also played a role. Therefore, we quantified the protein levels of four selected helicases (DDX23, DDX27, DDX42, and DDX50) using Western blot in MOCK and infected ZAP-WT and ZAP-KO cells (the same MOI and sampling time as for the ChIRP-MS experiment). DDX42 and DDX27 were chosen because they were among the top three enriched helicases (**Table S1I**). DDX23 and DDX50 are known to have antiviral activity [41,42].

Western blot analysis revealed expression of all four RNA helicases in both VERO-ZAP-KO and VERO-ZAP-WT cells, under mock condition and during Zika virus infection (**Fig 4A**). The expression of DDX23 and DDX27 was lower in both MOCK and Zika virus-infected ZAP-KO cells compared to ZAP-WT cells (DDX23 p =0.0374; p =0.0471; DDX27 p < 0.0001; p =0.0004), although ZAP-KO cells still showed prominent DDX23 and DDX27 expression (**Figs 4A, B**). Zika virus infection significantly reduced DDX27 expression in both cell lines (ZAP-KO p=0.415; ZAP-WT p=0.001) (**Fig 4A, B**). The expression of DDX42 was significantly lower in mock-infected ZAP-WT than in ZAP-KO cells (p = 0.0179). Zika virus infection considerably reduced DDX42 expression in both cell lines, but the comparative expression in ZAP-KO and ZAP-WT cells during infection was similar (p = 0.2139) (**Figs 4A, B**). The expression of DDX50 was lower in both MOCK and Zika virus-infected ZAP-KO cells compared to ZAP-WT cells (MOCK p = 0.0001; Zika p < 0.0001), although ZAP-KO cells still showed prominent DDX50 expression (**Figs 4A, B**). Zika virus infection did not affect DDX50 expression in either cell line (**Fig 4A, B**).

**Fig 4.**
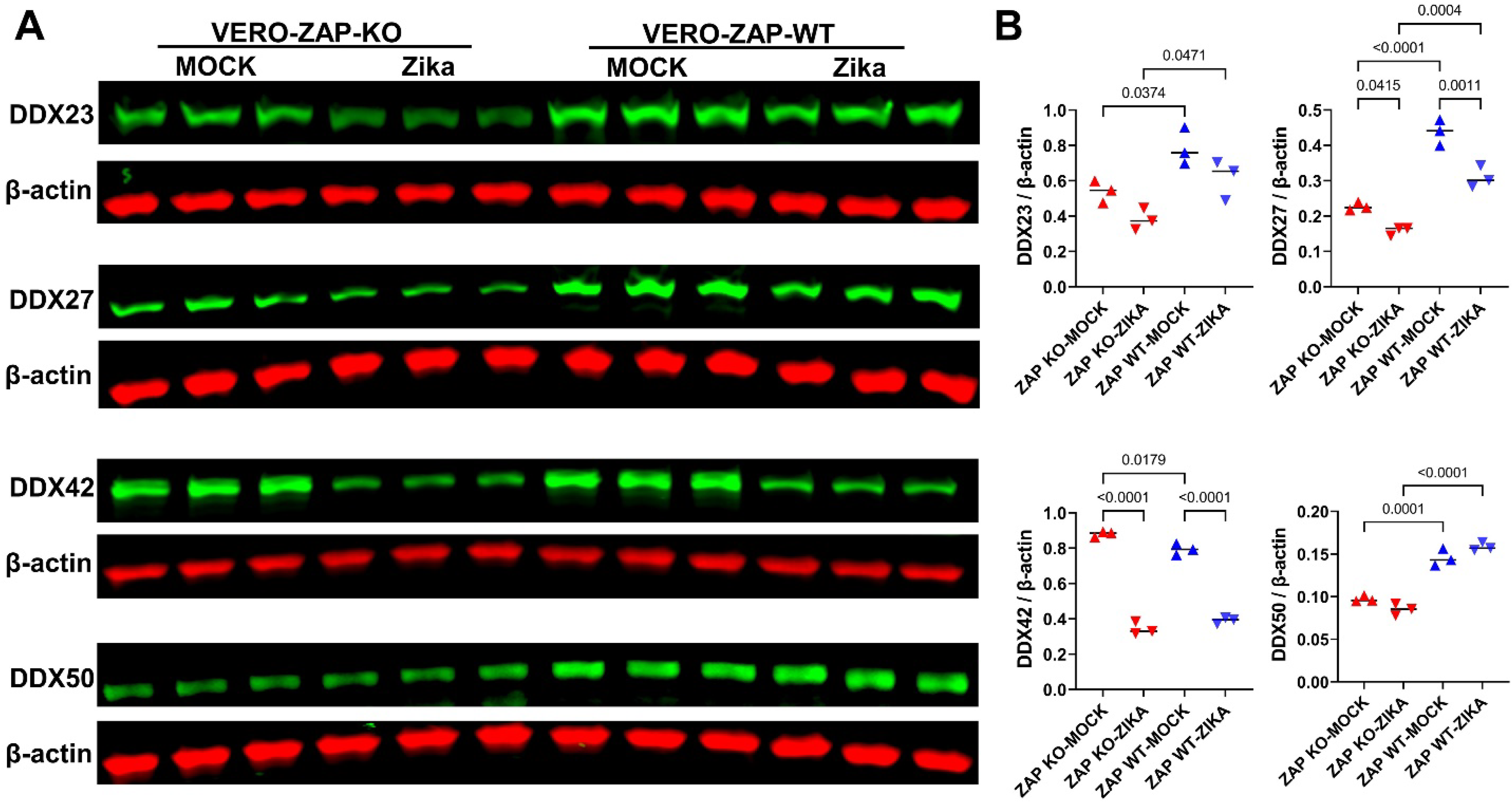
The expression of RNA helicases in VERO-ZAP-WT and VERO-ZAP-KO cells. Cells were mock infected or infected with normalized Zika virus MOIs; MOI 2 or 10 for VERO-ZAP-KO and VERO-ZAP-WT cells, respectively, as described for ChIRP in supplemental materials and methods. Cells were sampled at 72 h after mock or virus infection. (**A**) Green bands represent DDX23 (96 kDa), DDX27 (90 kDa), DDX42 (117-120 kDa), and DDX50 (83 kDa) RNA helicases. Red bands represent β-actin loading control (42 kDa). Western blot was done in 3 biological replicates for each experimental condition. (**B**) The expression of DDX RNA helicases was quantified as described in supplemental material and methods.

Overall, we confirmed that ZAP-WT and ZAP-KO cells express all four tested RNA helicases— DDX23, DDX27, DDX42, and DDX50. However, ZAP knockout was associated with reduced DDX23, DDX27, and DDX50 expression in both MOCK and infection conditions. These reduced cellular expressions of DDX23, DDX27, and DDX50 suggest that ZAP-mediated effects on cellular RNA helicase mRNAs may contribute to the depletion of RNA helicases in the ChIRP ZAP-independent Zika interactome. In contrast, DDX42 expression was even in ZAP-KO and ZAP-WT cells during Zika virus infection, indicating that direct interactions between ZAP, viral RNA, and cellular RNA helicases in ZAP-WT cells may play the central role in the enrichment of RNA helicases in the ChIRP ZAP-dependent Zika interactome.

## DISCUSSION

All previous ChIRP-MS studies on a viral RNA-cellular protein interactome were done in wild-type cell lines [8–11], and comparative ChIRP-MS studies in wild-type and KO cells were not reported. Here, using ChIRP-MS in ZAP-WT and ZAP-KO cells, we delineated ZAP-independent and ZAP-dependent cellular protein interactomes associated with Zika virus RNA. We showed that such experimental approach is feasible by normalizing MOIs and viral loads in ZAP-WT and ZAP-KO cells. Despite extensive research on ZAP’s antiviral activity, to our knowledge, this study is the first to explore the ZAP-dependent interactome specifically associated with viral RNA.

ZAP requires RNA helicases as co-factors: ZAP’s N-terminus interacts with the N– and C-terminal domains of DDX17 that promotes ZAP-mediated degradation of murine leukemia reporter virus [43]. However, our ChIRP-MS Zika data do not contain p72 DEAD-box RNA helicase (DDX17) (**Table S1B**). DHX30 interacts with ZAP, and the knockdown of DHX30 reduces ZAP’s antiviral activity against murine leukemia reporter virus [44]. Accordingly, we identified DHX30 in ChIRP-MS data (**Table S1B**); however, it was excluded from further interactome-specific analysis with our rigorous specificity cutoff. Interestingly, we found that 11 RNA helicases were enriched in the ZAP-dependent interactome—DDX42, DDX56, DDX27, DDX50, DDX52, DDX23, DDX54, DDX18, DHX57, DDX47, and one uncharacterized RNA helicase. In contrast, in the ZAP-independent interactome, no RNA helicases were enriched. The limitation is that our experimental design does not allow to identify the specific nature of interactions between Zika virus RNA and cellular proteins. We do not know yet which cellular proteins interact with only ZAP bound to viral RNA, which interact with ZAP and viral RNA, and which proteins have strict viral RNA dependency during interactions with ZAP. ZAP binds to single-stranded viral RNA and potentially to single overhangs in double-stranded RNA [45,46]; RNA helicases can bind to double-stranded RNA, single overhangs in double-stranded RNA, and single-stranded RNA; these make interactions between ZAP, viral RNA, and RNA helicases spatially possible. Additional research is necessary to determine if the RNA helicases identified in this study function as co-factors for ZAP in its antiviral activity, specifically ZAP-mediated degradation of viral RNA.

In addition to RNA helicases, ZAP requires other co-factors. It lacks intrinsic RNase activity but recruits the 5’ and 3’ RNA degradation machinery [24,47]. The E3 ubiquitin ligase TRIM25 is important for ZAP activity against alphaviruses by increasing inhibition of viral translation [48,49]. ZAP also interacts with the cytoplasmic protein KHNYN to inhibit CpG-enriched HIV-1 [50]. Riplet, a protein known to play a central role in activating the retinoic acid-inducible gene I (RIG-I), has been recently identified as a ZAP co-factor that augments the restriction of HIV-1 [51]. However, the full complexity of cellular proteins essential for ZAP antiviral activity is unknown. Large-scale interactome studies in non-infected cells have identified more than 250 proteins that may interact with ZAP [32,35–38]. Notably, there were no large-scale ZAP interactome studies in the context of viral infection [35]. Thus, our ZAP-dependent ChIRP-MS interactome (**Table S1H**) provides a highly specific list of potential ZAP co-factors, including eleven RNA helicases for future mechanistic and functional studies.

It is known that DDX RNA helicases not only facilitate viral RNA degradation but also act as viral RNA sensing proteins contributing to cellular innate immune signaling, type I IFN production, and antiviral response [52–57]. Also, viruses can abduct cellular RNA helicases to support their lifecycle and promote infectivity [58,59]. Cellular RNA helicases often interact with microbial genomes and evoke their properties in the complex with different adaptor proteins [52–54,56,60–62]. Thus, ZAP may play a dual role in mediating RNA helicase-dependent antiviral and proviral effects. Before our ChIRP-MS study, such co-factor/adaptor role of ZAP in interactions between viral RNA and cellular RNA helicases was not considered.

Interestingly, Western blot analysis showed that ZAP expression is associated with higher levels of at least three RNA helicases—DDX23, DDX27, and DDX50—since their expression was reduced in both MOCK and infected ZAP-KO cells as compared to ZAP-WT cells (**Fig 4**). Thus, ZAP may indirectly contribute to antiviral cellular responses by maintaining the expression of RNA helicases, which are known to have antiviral activity, including DDX23 and DDX50 [41,42].

Another interesting finding is that Zika virus infection considerably reduced the expression of DDX23, DDX 27, and DDX42 in ZAP-WT cells (**Fig 4**), as well as ZAP (**Fig S3**) under our experimental conditions (72 h post-infection). These observations may indicate Zika virus strategies for evading cellular immune responses, warranting further investigation.

The primary limitation of this study is its focus on an associative ChIRP-MS comparison between ZAP-WT and ZAP-KO cells. While conducting functional studies could provide further insights, it falls beyond the scope of this work. Nonetheless, our ZAP-dependent interactome offers the scientific community a valuable resource for future studies on ZAP, its co-factors, and viral RNA interactions. It would also be interesting to compare ChIRP-MS interactomes across different ZAP-WT and ZAP-KO cell lines infected with various viruses. Our strict filtering approach to identify the ZAP-dependent interactome, which excluded proteins from the ZAP-independent interactome even with a single MS spectral count in ZAP-KO cells, might be seen as too rigorous. However, it ensures a highly specific list of potential ZAP co-factors. Additionally, we have made available the raw and extended ZAP interactome databases (**Table S1B, C; Supplemental Dataset 2**), allowing others to apply their own filtering approaches.

## CONCLUSIONS

Together, the ZAP-dependent interactome identified with ChIRP-MS provides potential ZAP co-factors for antiviral activity against Zika virus and possibly other viruses. Identifying the full spectrum of ZAP co-factors and mechanisms of how they act will be critical to understand the ZAP antiviral system and may contribute to the development of antivirals.

## MATERIALS AND METHODS

Details of ChIRP-MS, cells, viruses, comparative infection studies, RT-qPCR, Western blot, and statistics are in Supplemental Materials and Methods.

## AUTHOR CONTRIBUTIONS

Conceptualization: UK. Investigation: NPKL, AJS, PPS, UK. Data analysis: NPKL, AJS, PPS, UK. Funding: UK. Resources: UK. Writing—original draft preparation: NPKL, AJS, PPS, UK. Writing—review and editing: UK, NPKL, PPS.

## DATA AVAILABILITY

Data available within the article, its supplementary materials, and DRYAD open data publishing platform which can be accessed via links: https://datadryad.org/stash/share/tqLeKFRU68c947MAFplLF9S8xrd_JM8IulT5JFSiEio and https://datadryad.org/stash/share/GJWy9A9Hofp-9aya9oOLI9RgDsi_qqw6FITwPu4F0Gs

## SUPPORTING INFORMATION CAPTIONS

### Supplemental Materials and Methods

The file contains experimental details.

**Fig S1.**
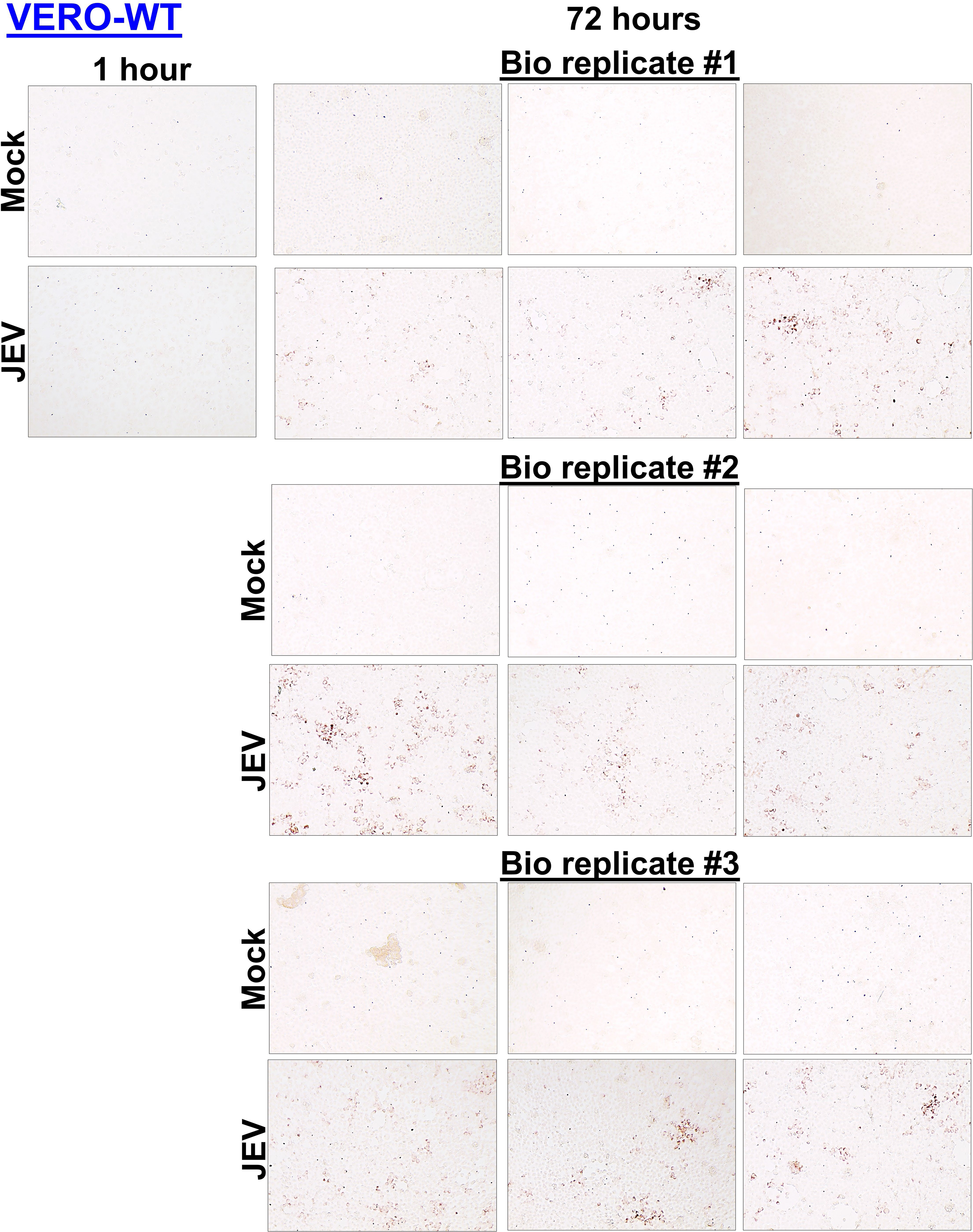
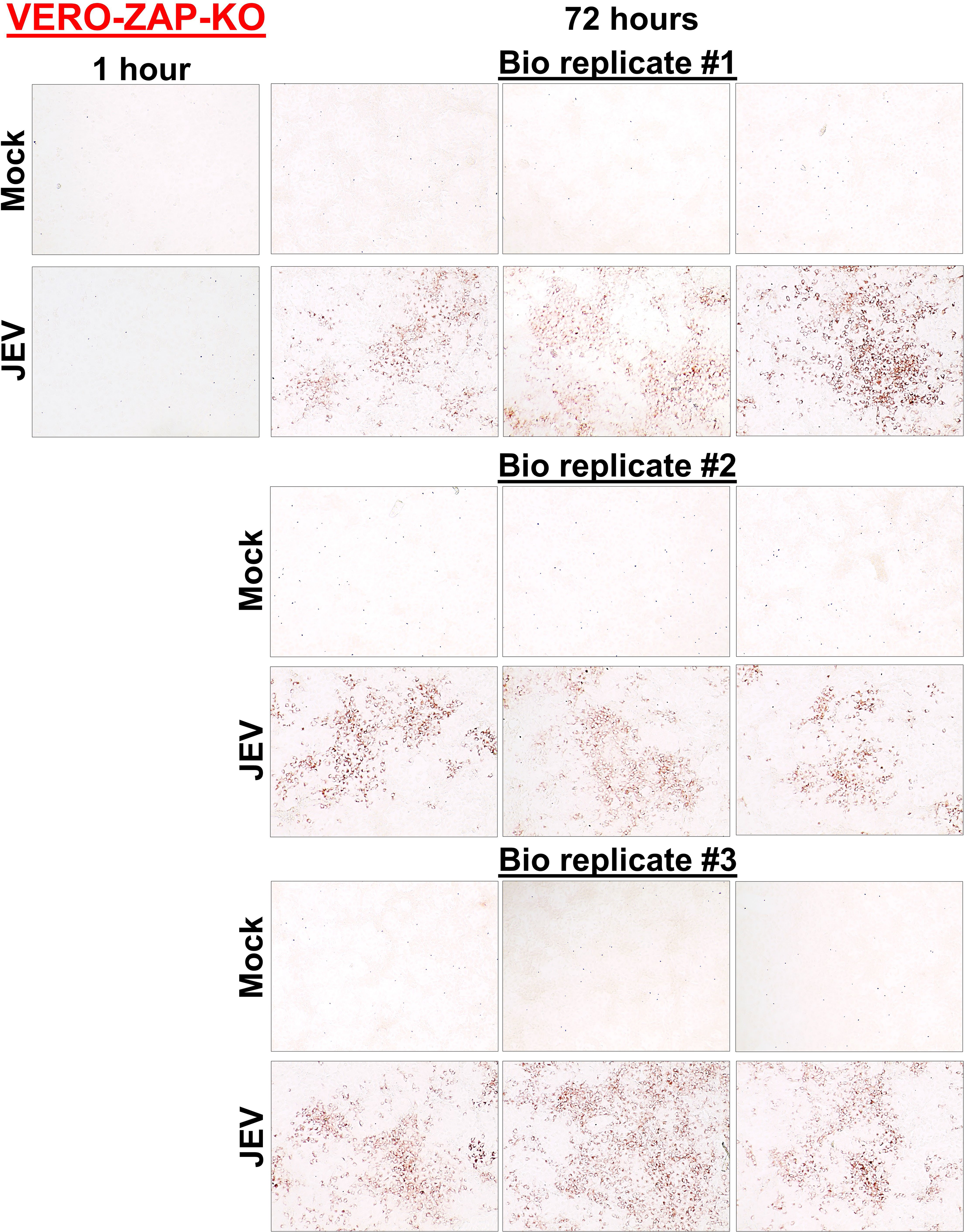
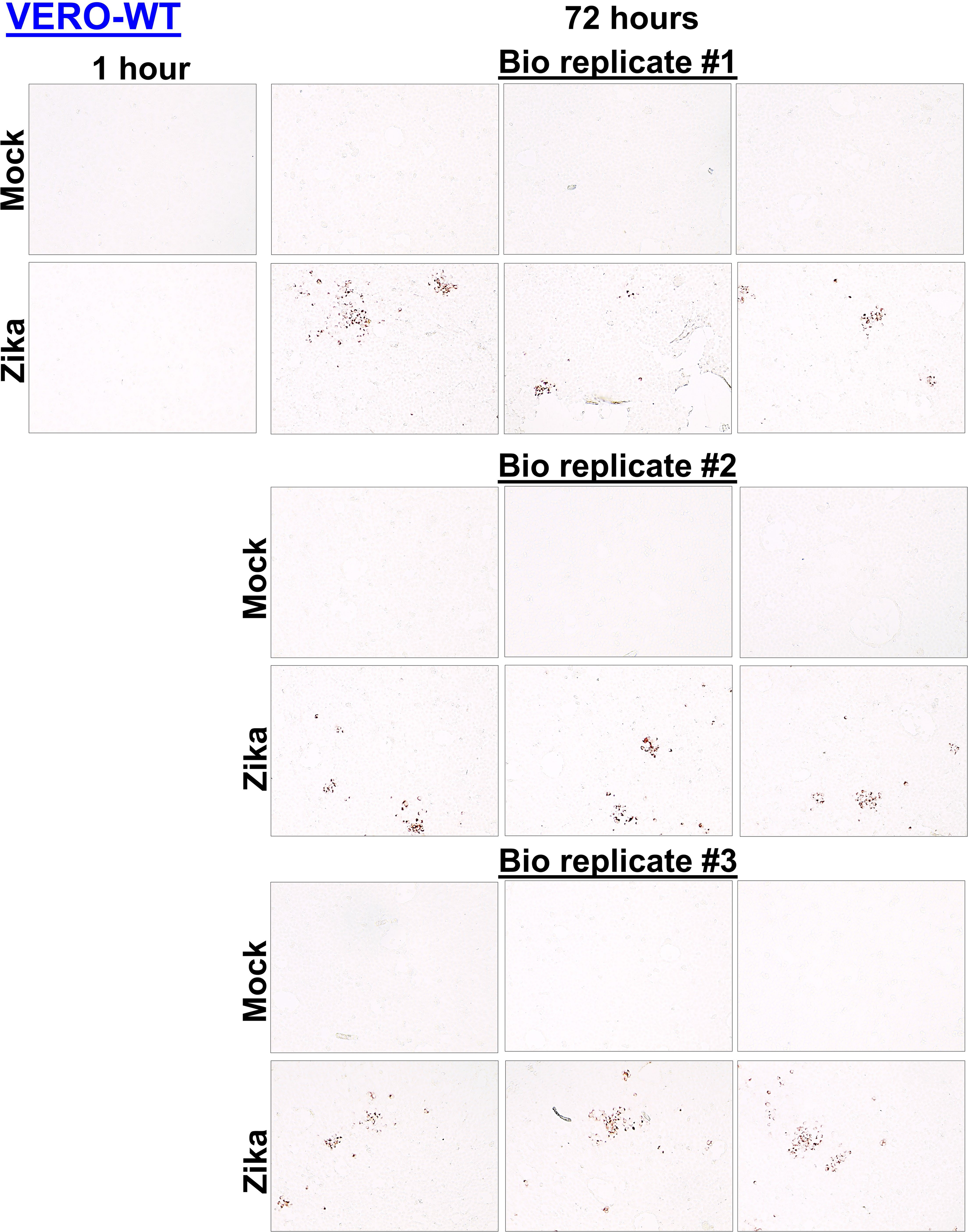
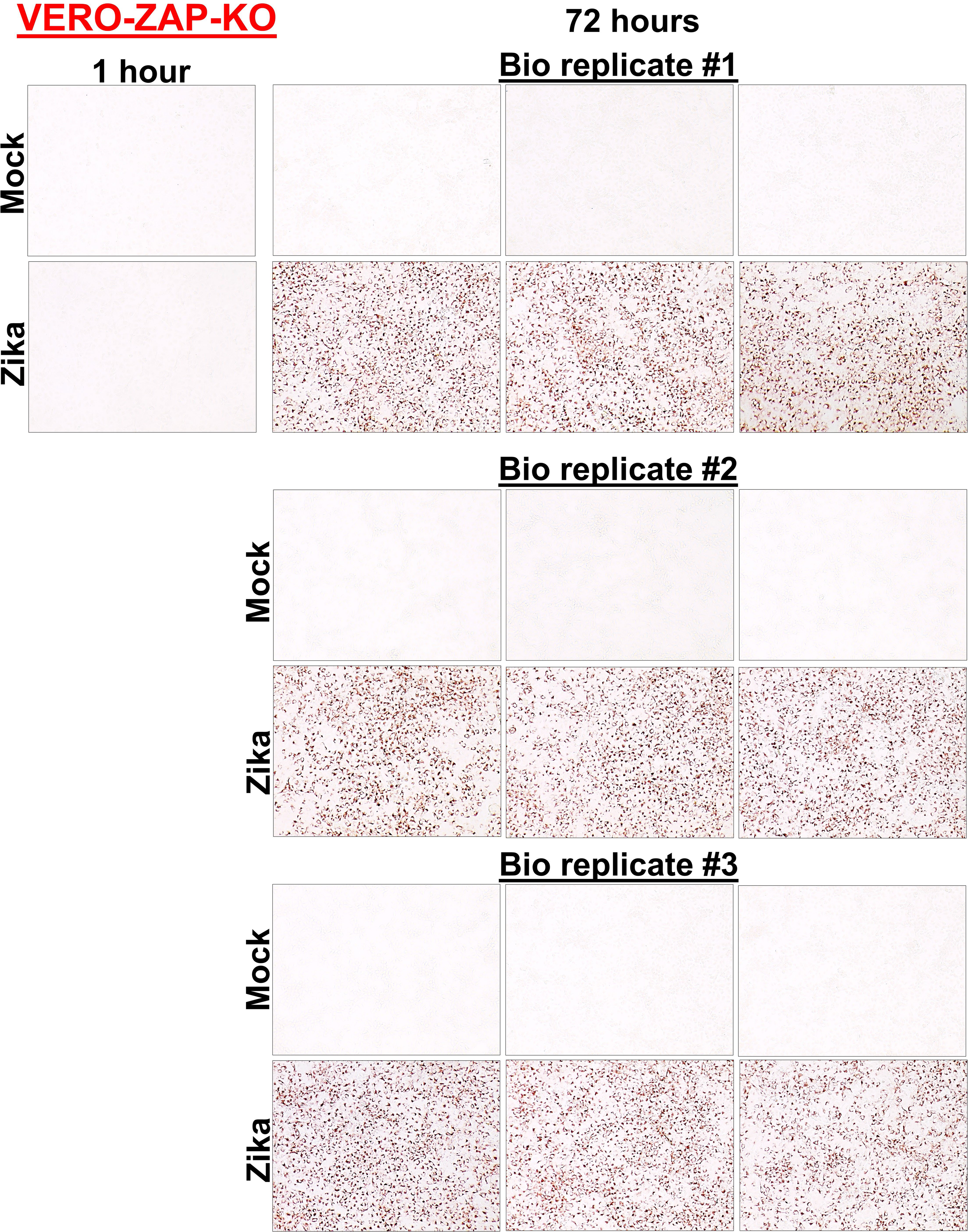
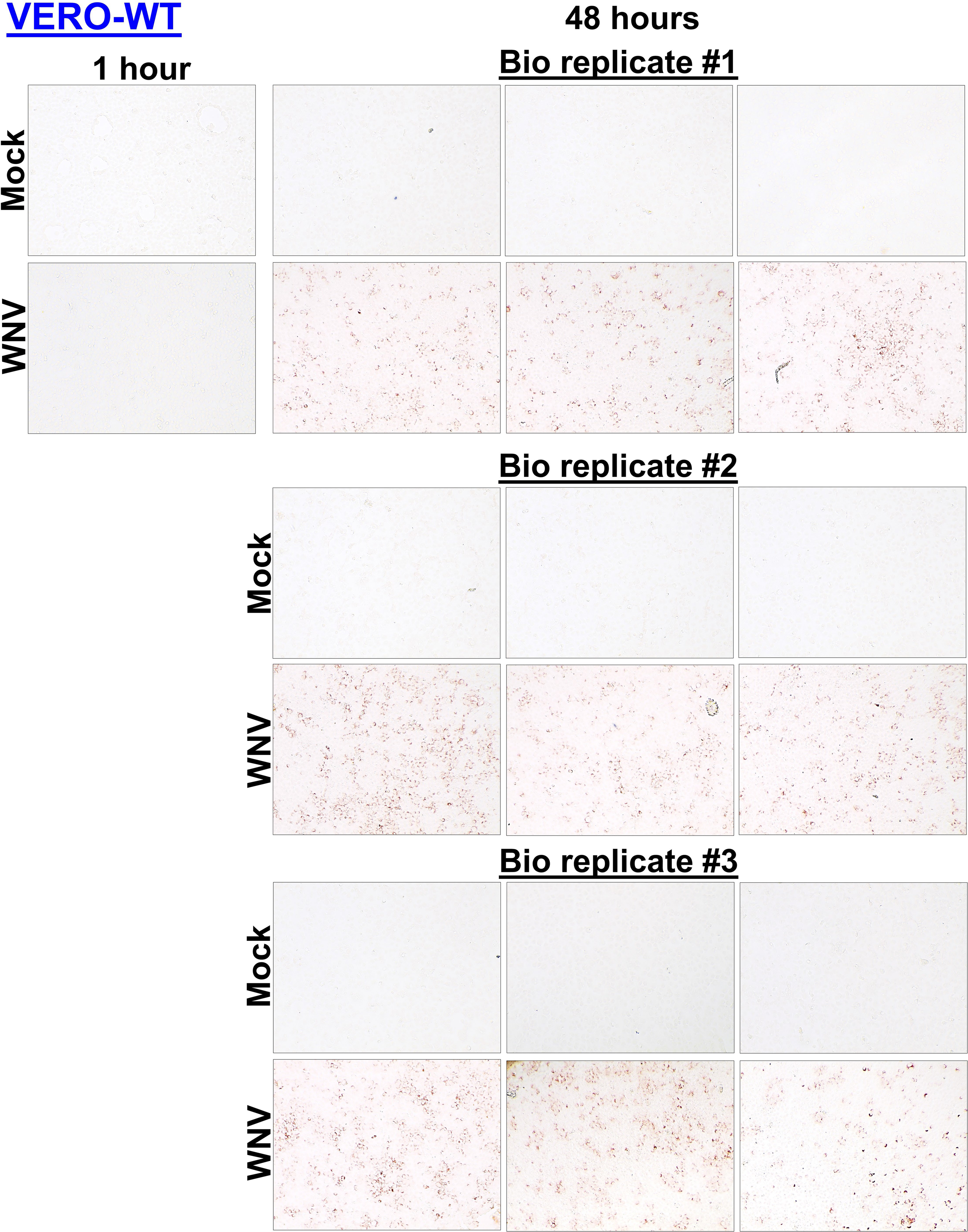
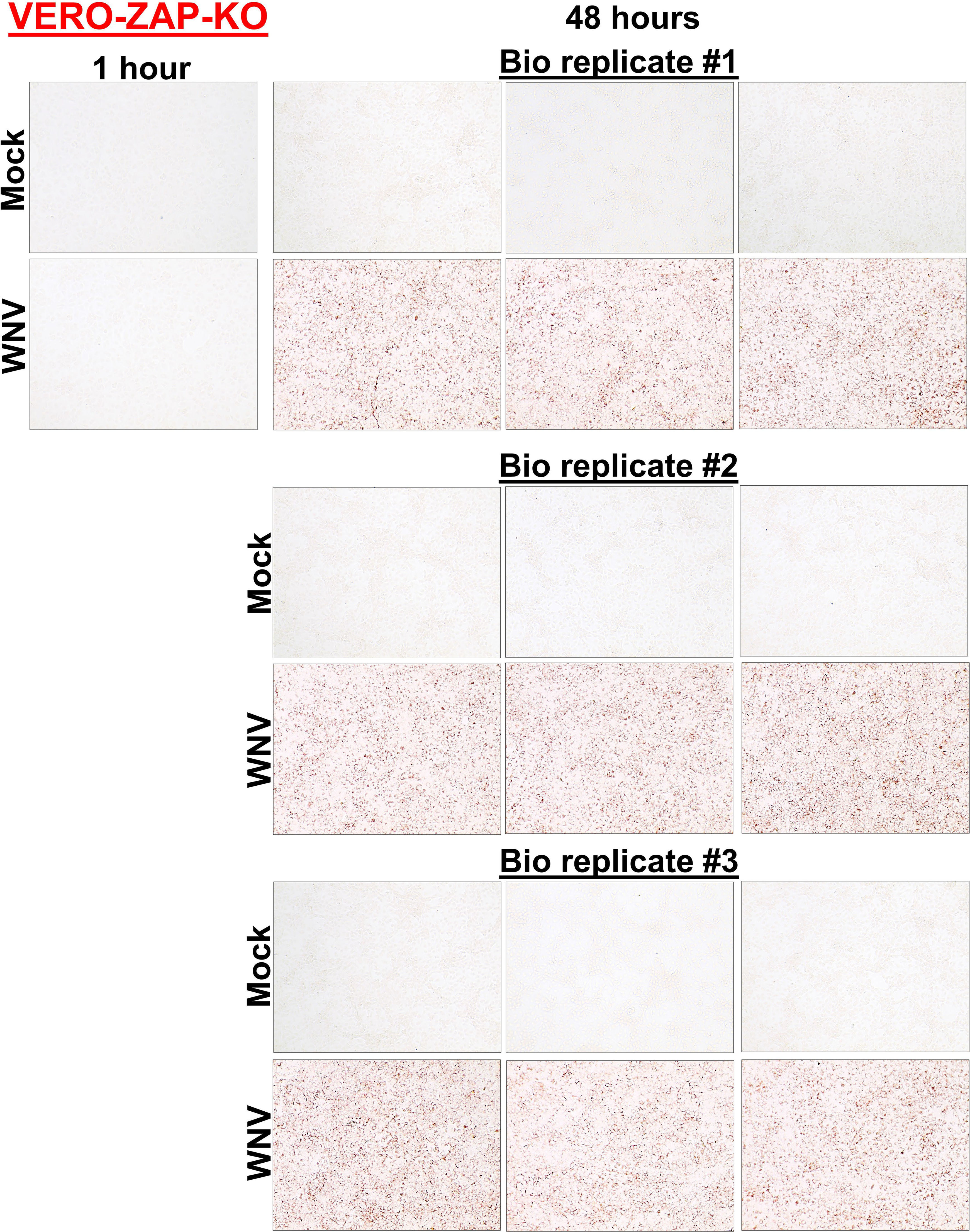
Representative images of cells positive for flavivirus E protein (red staining) at 72 h (JEV and Zika virus) and 48 h (WMV). The experiment was done in 3 biological and 3 technical replicates. All replicates are shown. MOI 0.01. The data from these replicate figures were used for digital quantification in **Figs 1B**, **F**, **J**. (**A**) JEV in VERO-ZAP-WT cells. (**B**) JEV in VERO-ZAP-KO cells. (**C**) Zika virus in VERO-ZAP-WT cells. (**D**) Zika virus in VERO-ZAP-KO cells. (**E**) WNV in VERO-ZAP-WT cells. (**F**) WNV in VERO-ZAP-KO cells.

**Fig S2.**
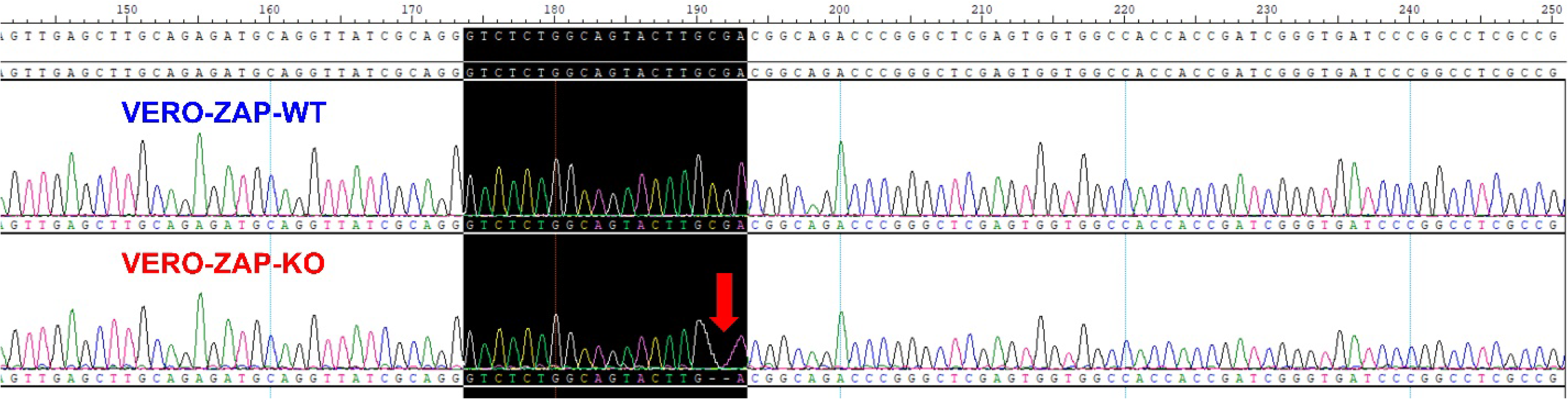
VERO-ZAP-WT and VERO-ZAP-KO cells genotyped with Sanger sequencing as described in Supplemental Materials and Methods. Location of indels on *Chlorocebus sabaeus* genome: –2bp deletion, NW_023666072.1: 4677550-4677551.

**Fig S3.**
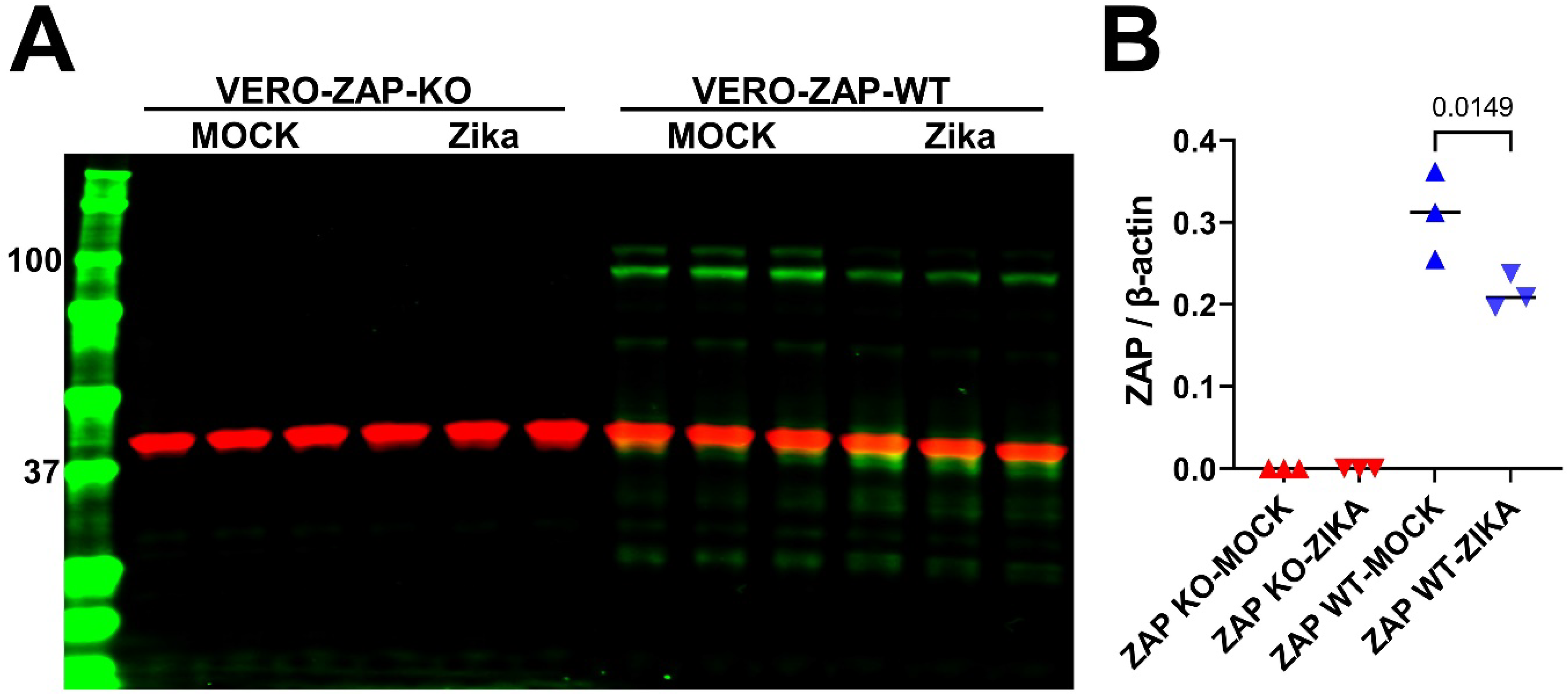
ZAP Western blot. (**A**) ZAP Western blot in VERO-ZAP-KO and VERO-ZAP-WT cells. Cells were mock infected or infected with normalized Zika virus MOIs; MOI 2 or 10 for VERO-ZAP-KO and VERO-ZAP-WT cells, respectively, as described for ChIRP in supplemental materials and methods. Cells were sampled at 72 h after mock or virus infection. Green bands represent ZAP with the manufacturer’s predicted size 111 kDa; multiple bands may represent four isoforms described for human ZAP [31] and still not experimentally characterized for monkeys. Red bands represent β-actin loading control, 42 kDa. Western blot was done in 3 biological replicates for each experimental condition. (**B**) The expression of ZAP was quantified as described in supplemental material and methods.

**Fig S4.**
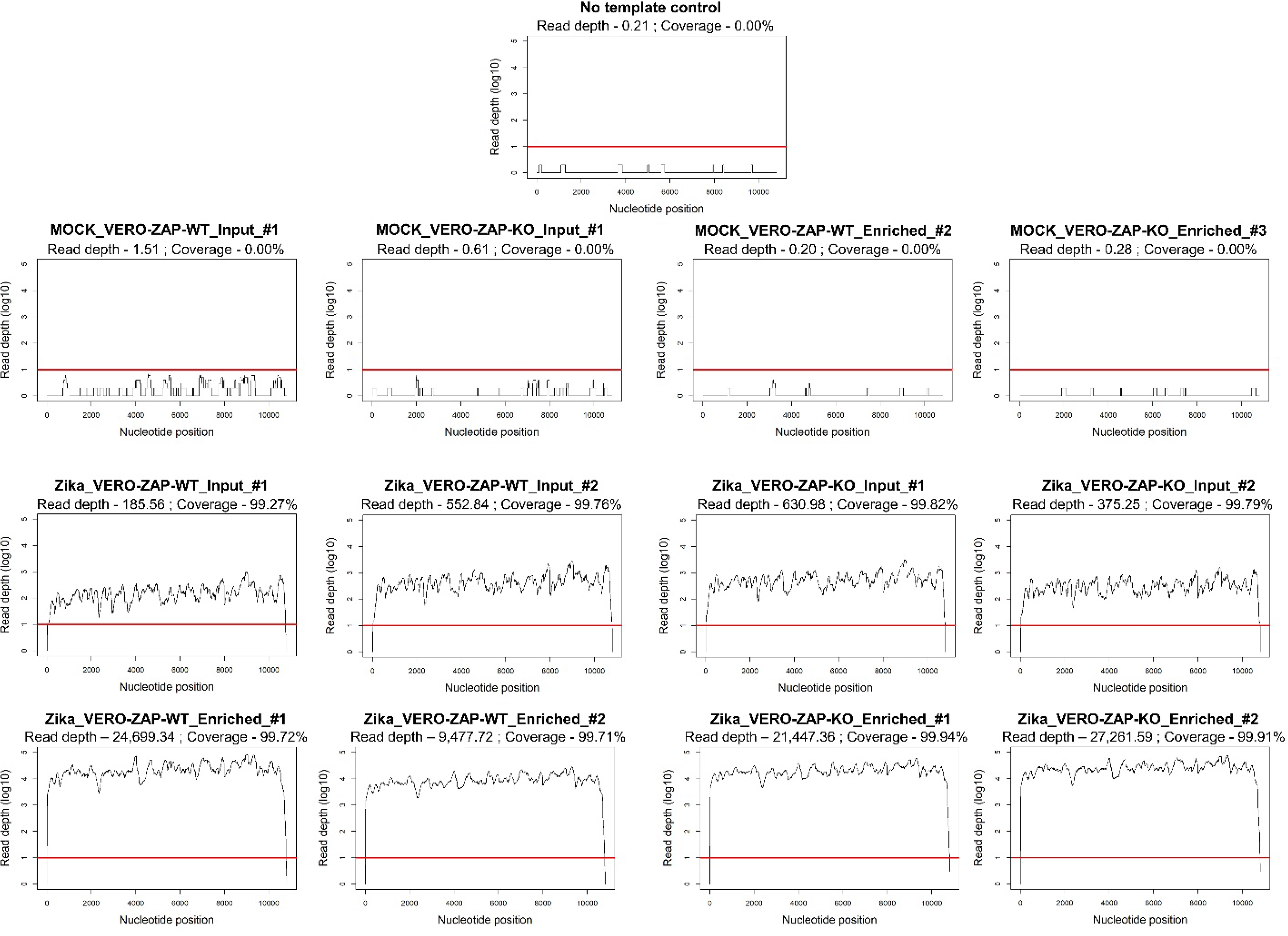
The Zika RNA genome coverage and NGS depth in VERO-ZAP-WT and VERO-ZAP-KO cells. **Input** – sonicated cellular lysates. **Enriched** – immunoprecipitated viral RNA-protein complexes bound to MyOne C1 magnetic beads at the last washing step before protein elution. Control NGS in mock-infected cells did not show Zika-specific sequences. A red line shows the 10-nucleotide NGS depth threshold.

**Table S1.** – ChIRP-MSdata. The table contains raw and analyzed ChIRP-MS data. The table can be accessed in the DRYAD open data publishing platform: https://datadryad.org/stash/share/tqLeKFRU68c947MAFplLF9S8xrd_JM8IulT5JFSiEio

**Supplemental Dataset 1.** This dataset contains 26 raw fastq files representing paired-end NGS data for **input** and **enriched** samples described in results, in **Figure 2D**, and in **Fig S4**. The dataset also contains the “Zika virus NGS sample IDs” file which provides identifications for raw files and experimental conditions (VERO-ZAP-WT or VERO-ZAP-KO cells; MOCK or Zika virus; input or enriched samples; replicate number). The dataset can be accessed in the DRYAD open data publishing platform: https://datadryad.org/stash/share/GJWy9A9Hofp-9aya9oOLI9RgDsi_qqw6FITwPu4F0Gs

**Supplemental Dataset 2.** This dataset contains the folder “RAWfiles” with 12 Mass spectrometry RAW data files representing three biological replicates for MOCK and Zika virus-infected VERO-ZAP-WT and VERO-ZAP-KO cells. The folder “MSFragger” contains the output of FragPipe v18.0 which was used to analyze Mass spectrometry RAW data files as described in Supplemental Materials and Methods. The folder “MSFragger” contains data for individual samples, and the “combined_protein.tsv” file (also represented in **Table S1B**), which was used for the identification of Extended, ZAP-independent, and ZAP-dependent Zika virus RNA interactomes from ChIRP-MS results as described in Supplemental Materials and Methods. The “Zika virus ChIRP-MS sample IDs” file provides identifications for Mass spectrometry and FragPipe raw files and experimental conditions (VERO-ZAP-WT or VERO-ZAP-KO cells; MOCK or Zika virus; biological replicate number). The dataset can be accessed in the DRYAD open data publishing platform: https://datadryad.org/stash/share/GJWy9A9Hofp-9aya9oOLI9RgDsi_qqw6FITwPu4F0Gs

## SUPPLEMENTAL MATERIALS AND METHODS

### Cells

Wild-type and ZAP knockout VERO cells were previously described [1]. Briefly, wild-type VERO E6 cells (ATCC) (VERO-ZAP-WT) and ZAP knockout (VERO-ZAP-KO) derivatives were maintained in DMEM (Fisher, MA, USA; #11-965-118) supplemented with 10% heat-inactivated fetal bovine serum (FBS) (Fisher, MA, USA; #A5256801), 1x Penicillin-Streptomycin (Fisher, MA, USA; #15140122), and 2.67 mM sodium bicarbonate (Fisher, MA, USA; # 25080094) at +37°C in a 5% CO_2_ humidified incubator. To generate Zinc finger CCCH-type antiviral protein 1 (ZAP) KO cell line, we used the guide RNA (gRNA) GTCTCTGGCAGTACTTGCGA which targets the first exon of *Chlorocebus sabaeus* (Gene ID: 103226990) ZC3HAV1 gene, which is required for the expression of all ZAP isoforms. A non-targeting control gRNA ACGGAGGCTAAGCGTCGCAA was also used. These gRNAs were transiently transfected into cells using GenCrispr NLS-Cas9-NLS Nuclease (GenScript). Transfected cells were plated in 96-well plates by limiting dilution to generate isogenic single clones. The clones were expanded and genotyped with Sanger sequencing; primers (F-ATCGCTGGGCTGGACTAACG, R-GCAGAGAAGGGAGTGGCTGAA) were used to identify indels (location of indels on *Chlorocebus sabaeus* genome: –2bp deletion, NW_023666072.1: 4677550-4677551; **Fig S2**). The knock-out subclone verified by genotyping was further confirmed by western blot [1] (**Fig S3**). The negative control cells were not isolated into single clones, but they were validated by Sanger sequencing to confirm there were no indels (**Fig S2**). Free mycoplasma contamination status in cells was confirmed using the LookOut Mycoplasma PCR Detection Kit (Millipore Sigma, MA, USA; #MP0035).

### Viruses

We used the Asian lineage Zika virus H/PF/2013 strain. Zika virus was synthetically rescued with reverse genetics in our previous study [2]; H/PF/2013 [GenBank: KJ776791.2] reference sequence was used to rescue the virus. For JEV (SA14-14-2 strain, GenBank MK585066.1) and WNV (NY99 strain, GenBank: DQ211652.1), we used the same reverse genetics system [2]. Viruses were propagated in VERO cells in DMEM supplemented with 2% FBS and 1% penicillin/streptomycin (P/S). Supernatants were centrifuged (12,000 g, 20 min, +4°C), aliquoted, and frozen (–80°C). Infectious virus titers were quantified using VERO E6 cells with the endpoint dilution assay, and fifty percent tissue culture infective dose (TCID_50_) endpoint titers were calculated as previously described [2–5]. Media from virus-negative VERO cells was processed in the same manner and used in control conditions. Free mycoplasma contamination status in virus stocks was confirmed using the LookOut Mycoplasma PCR Detection Kit (Millipore Sigma, MA, USA; #MP0035).

### Comparative infection phenotypes in wild-type and ZAP-KO cells infected with Zika virus, JEV, and WNV

We tested alongside JEV, Zika virus, and WNV in VERO-ZAP-WT and VERO-ZAP-KO cells. Fifty thousand cells per well were seeded into 96-well plates; next day, media was removed, and monolayers were inoculated with MOIs 0.01 of JEV, Zika, or WNV in 50 µl for 2 h at +37°C and 5% CO_2_. Afterward, the inoculum was removed and replaced with 150 µl DMEM supplemented with 2% FBS, 1x Penicillin-Streptomycin, and 2.67 mM sodium bicarbonate. At 1 h and 72 h (48 h for WNV), cells in 9 wells representing three biological and three technical replicates were fixed and stained for flavivirus E protein as we did [2,5–9]. For immunohistochemistry staining we used 4G2 (ATCC, MA, USA; #HB-112) antibodies (Ab) against Zika and WNV E protein, and #GTX125867 Ab (GeneTex, CA, USA) against JEV E protein [2–5]. To quantify and compare immunohistochemistry staining between ZAP-WT and ZAP-KO conditions, a magnification x200 images were randomly acquired in all replicates with the same microscopic settings. Afterward each image was procced in ImageJ (NIH, USA) with Automatic Particle counting (https://imagej.net/imaging/particle-analysis). In addition, cell lysates from 3 biological and 2 technical replicates were collected at the same time points with 140 µl of PureLink RNA Mini Kit lyses buffer (Invitrogen) for subsequent RNA extraction and virus-specific RT-qPCR [4,5,8,10]. The difference between ZAP-WT and ZAP-KO conditions was represented by fold change of viral RNA loads in cell lysates collected at 1 h and 72 h (48 h for WNV) after inoculation (1 h after inoculation was used to normalize for leftover virus inoculum RNA).

### RNA extraction and virus-specific reverse transcriptase quantitative polymerase chain reaction assays (RT-qPCR)

To quantify intracellular Zika virus, JEV, and WNV RNA, total RNA was extracted from cell lysates with PureLink RNA Mini Kit (Invitrogen) according to the manufacturer’s instructions. All RT-qPCR reactions were conducted on the Applied Biosystems QuantStudio 3 real-time PCR system (Fisher, MA, USA) and analyzed using QuantStudio Design & Analysis Software v1.5.2.

For Zika virus RNA quantification, we used the previously validated SYBR one-step RT-qPCR assay [11]. The Luna Universal One-Step RT-qPCR Kit (NEB, MA, USA; # E3005L) reaction mixture (20 μL) consisted of 10 μL Luna Universal One-Step Reaction Mix, 1 μL Luna WarmStart RT Enzyme Mix, 1 μL (10 μM) of forward (ZIKV-F-SYBR: 5′-AGGATCATAGGTGATGAAGAAAAGT-3′) and reverse (ZIKV−R-SYBR: 5′-CCTGACAACACTAAGATTGGTGC-3′) primers, 3.4 μL nuclease-free water and 4 μL of RNA template. The reverse transcription and enzyme activation steps of 10 min at 55°C and 1 min at 95°C were followed by 40 amplification cycles (10 s at 95°C and 60 s at 60°C). A standard curve was generated and used to quantify Zika RNA loads as we described [1,2,4].

For JEV (F: 5′-GCCACCCAGGAGGTCCTT-3′; R: 5′-CCCCAAAACCGCAGGAAT-3′; Probe: 56-FAM-CAAGAGGTG/ZEN/GACGGCC-3IABkFQ) and WNV (F: 5′-AGTAGTTCGCCTGTGTGAGC-3′; R: 5′-GCCCTCCTGGTTTCTTAGA-3′; Probe: 5’-FAM-AATCCTCACAAACACTACTAAGTTTGTCA-TAMRA-3’), we used the previously validated probe-based one-step RT-qPCR assays [12,13]. The Luna Universal Probe One-Step RT-qPCR Kit reaction mixture (20 μL) consisted of 10 μL Luna Universal Probe One-Step Reaction Mix, 1 μL Luna WarmStart RT Enzyme Mix, 1 μL (10 μM) of forward and reverse primers, 0.5 μL of the probe, 2.5 μL nuclease-free water and 4 μL of RNA template. The reverse transcription and enzyme activation steps of 10 min at 55°C and 1 min at 95°C were followed by 40 amplification cycles (10 s at 95°C and 30 s at 60°C). A standard curve was used to quantify viral RNA loads as we described [1,2,4,6].

PCR values were corrected for volumes (140 µl of cell lyses buffer) and upon logarithmical transformation expressed as virus RNA genome copies per mL. Strict precautions were taken to prevent PCR contamination. All master mix preparations were done in the dedicated PCR cabinet. Aerosol-resistant filter pipette tips and disposable gloves were used. Kit reagent controls were included in every RNA extraction and PCR run.

### Western blot

Cells in T-75 or T-175 flasks were washed, pelletized, and lysed in 150 or 300 μL of RIPA buffer (Fisher, MA, USA; # PI89900) supplemented with 1× Halt protease inhibitor cocktail (Fisher, MA, USA; # PI87786) and 1× EDTA (Fisher, MA, USA; #BP2482100). After 10 minutes of incubation on wet ice, the suspension was gently vortexed and centrifuged (12,000g, 10 min, +4 °C). Supernatants were aliquoted in prechilled low protein binding tubes and stored at −80°C. After quantifying protein concentration with the Pierce BCA Protein Assay Kit (Fisher, MA, USA; # PI23225), 50 μg of protein extract was mixed with 4x reducing Laemmli SDS sample buffer (Fisher, MA, USA; # AAJ60015AC), boiled at 95°C for 10 min, and resolved using a 10% Mini-PROTEAN TGX precast protein gels (Bio-Rad, CA, USA; #4561036) running in 1X Tris/Glycine/SDS running buffer (Bio-Rad, CA, USA; # 1610772) at a constant current of 125V for 70 min. Proteins were transferred to methanol-activated 0.45 μm Immun-Blot Low Fluorescence polyvinylidene fluoride (PVDF) (Bio-Rad, CA, USA; #1620260) membranes using Trans-Blot Turbo RTA Mini 0.45µm LF PVDF Transfer Kit (Bio-Rad, CA, USA; #1704274). Afterward, PVDF membranes were blocked with 1X VWR Life Science Rapidblock Blocking Solution (VWR, GA, USA; #97064-124) for 1 h at room temperature. The blocked membranes were incubated with primary antibody diluted in blocking solution overnight at +4°C. Membranes were washed four times with TBST (1x Bio-Rad Tris Buffered Saline containing 0.1% of Fisher Tween-20). After that, membranes were incubated with secondary antibody along with hFAB™ Rhodamine Anti-Actin Primary Antibody (Recombinant human β-actin) diluted in Blocking Solution for 1 h at room temperature, and washed four times with TBST. Fluorescent blots were imaged on the ChemiDoc MP Imaging system (Bio-Rad, CA, USA) using an appropriate detection channel with Image Lab Touch Software (Bio-Rad, CA, USA).

We used antibodies (Abs): Rabbit anti-ZAP IgG Abs (1:1,000 dilution; ANTI-ZC3HAV1 #HPA059096-100UL, Millipore Sigma, MA, USA), Rabbit anti-DDX23 IgG Abs (1:1,000 dilution; #ab70461, Abcam, MA, USA), Rabbit anti-DDX27 IgG Abs (1:1,000 dilution; #ab177938, Abcam, MA, USA), Rabbit anti-DDX42 IgG Abs (1:2,000 dilution; Fisher, MA, USA; #A303-353A), Rabbit anti-DDX50 IgG Abs (1:1,000 dilution; #10358-1-AP; ProteinTech, IL, USA), Rabbit anti-Zika virus NS5 IgG Abs (1:1,000 dilution; #GTX133312; GeneTex, CA, USA), Rabbit anti-West Nile virus NS5 IgG Abs (1:1,500, #GTX131961; GeneTex, CA, USA), Rabbit anti-Japanese Encephalitis virus NS5 Abs (1:1,500, #GTX131359; GeneTex, CA, USA), hFAB™ Rhodamine Anti-Actin Primary Abs (1:3,000; #12004164; Bio-Rad, CA, USA), and IRDye® 680RD Goat anti-Rabbit IgG secondary Abs (1:10,000; #926-68071; LI-COR, NE, USA).

Image Lab Software Version 6.1 (Bio-Rad, CA, USA) was used for Western blots densitometry analysis according to the user guide instructions. Briefly, the “Lanes and Bands” option was used to select the specific lanes and bands of the corresponding DDX RNA helicase, ZAP, and β-actin lanes and bands. The “Lane Profile” option was used to subtract the background. In the “Analysis Table,” the background-adjusted (Adj. Volume) intensities of each DDX RNA helicase and ZAP bands were normalized by the background-adjusted β-actin intensities. Subsequently, the DDX RNA helicase (or ZAP)/β-actin ratios were used to analyze data for each experimental condition.

### Comprehensive Identification of RNA binding Proteins by Mass Spectrometry (ChIRP-MS)

We performed ChIRP-MS as described before [14–17]. Biotinylated DNA probes (**Table S2A**) covering the whole Zika virus genome were designed using the online tool (https://www.biosearchtech.com/chirp-designer). To have an even coverage of the whole genome, 50 probes were designed by dividing the whole Zika reference sequence H/PF/2013 [GenBank: KJ776791.2] into two segments (1-5400 and 5401-10807), with a repeat masking setting of 3 and spacing length of 80 (1 oligo around every 200 bp). Oligos were synthesized with 3’ biotin-TEG modification, final yield of 2 OD, and HPLC purification at Bio Basic Inc, Canada. Fifty individual oligos were resuspended at 100 µM and mixed at equal volumes to prepare the probe pool.

For Zika virus infection, VERO-ZAP-WT and VERO-ZAP-KO cells were seeded at 10^7^ cells per T-175 flask (4 flasks per experimental condition x 3 biological replicates) and were grown in DMEM supplemented with 10% FBS. The next day, the media was removed, and cell monolayers were washed with 1× PBS and inoculated with Zika virus at MOI 10 for VERO-ZAP-WT cells and MOI 2 for VERO-ZAP-KO cells; these MOIs were adapted to provide the same Zika virus loads. For this, we incrementally increased the MOI for VERO-ZAP-WT cells, as Zika virus caused a more aggressive infection in VERO-ZAP-KO cells (**Figs 1E-H**). To exclude the effects of differential uptake of the stimulus with MOI 10 in VERO-ZAP-WT cells, during VERO-ZAP-KO MOI 2 Zika virus inoculation we added heat-inactivated virus equivalent of MOI 8. Control cells were mock-treated with supernatant from virus-free control cells. Flasks were incubated at 37°C for 2 h and gently shaken every 10 minutes. The inoculum from flasks was removed, and monolayers were extensively washed with 1× PBS twice and 25 mL of DMEM supplemented with 2% FBS was added to each flask.

After 72 h, for cell harvesting and chemical crosslinking, media was removed, cell monolayers were rinsed with 1× PBS, trypsinized, washed, and cells from four T-175 flasks representing one biological replicate were pooled for each experimental condition. Cells were cross-linked in 37 mL of 3% methanol free-formaldehyde (Fisher, MA, USA; #PI28908) diluted in 1× PBS and incubated for 30 min at room temperature on a rotator. Afterwards, formaldehyde was quenched with 3.7 mL of 1.25 M glycine (Fisher, MA, USA; #AAA1381636) with final concentration 125 mM and pelleted (2,000g, 5 min, +4°C). Cell pellets were washed, resuspended in 2.5ml of 1× PBS, transferred to pre-weighed microcentrifuge tubes, and centrifuged (10,000g, 2 min, +4°C). PBS was aspirated, tubes were placed on ice and weighted on an analytical scale to determine the final pellet weight, snap frozen in liquid nitrogen, and stored at –80°C. Around 300 mg of cells was obtained from four combined T-175 flasks (4 flasks per experimental condition x 3 biological replicates).

For cell lysis and sonication, cell pellets were thawed at RT, resuspended in the ChIRP lysis buffer (50 mM Tris-HCl, pH 7.0, # BP1756500; 10 mM EDTA, #AAJ62786AK; 1% SDS, # BP2436-200; all reagent are from Fisher, MA, USA) supplemented just before use with PMSF (100 mM; #PI36978, Fisher, MA, USA; 10 µl per 1 ml of ChIRP lysis buffer), SUPERaseIn (20 U/ µl; #AM2694, Fisher, MA, USA; 10 ul per 1 ml of ChIRP lysis buffer), and protease inhibitor (100x; #P178410, Fisher, MA, USA; 10 ul per 1 ml of ChIRP lysis buffer) to finally have 300 mg of cell pellet per 4 ml of the complete lysis buffer. The lysate from each sample were aliquoted in 2 ml and sonicated using Sonics Ultrasonic Processor VCX750 with 3 mm microtip at 20% amplitude with 15 seconds On, 45 seconds Off pulse intervals for 1 h (1 h for total 15 + 45 seconds On and Off intervals) in 15 mL MI Bioruptor Plus TPX Tubes (#C30010009; HOLOGIC Diagenode, NJ, USA). The sonicated lysates were snap frozen in liquid nitrogen and stored at –80°C until further use. Aliquots of sonicated lysates (40 µL) were used to extract RNA/DNA with the Invitrogen PureLink RNA Mini Kit (#12-183-018A; Fisher, MA, USA) according to manufacturer’s instructions, and the efficiency of sonication was confirmed by running extracts from all samples on 1% agarose gels. The average DNA/RNA length was around 500 nucleotides in all samples. The RNA from these samples were also used for Zika-specific RT-qPCR and Next Generation Sequencing (NGS) to analyze the entire Zika virus RNA loads, genome coverage and NGS depth (see below and results).

For hybridization, sonicated lysates were thawed on ice and pre-cleaned by adding 30 μL washed Invitrogen Dynabeads MyOne Streptavidin C1 (# 65001; Fisher, MA, USA) per 1 mL of lysate and rotating at 37°C for 30 min in a hybridization oven. Beads were removed twice from lysates using a magnetic stand; for this and all subsequent magnetic stand steps > 2 min magnetization was applied before removing the supernatant. Next, for every 1 mL of pre-cleaned sonicated lysate, 2 mL of ChIRP hybridization buffer (750 mM NaCl, 1% SDS, 50 mM Tris-HCl pH 7.0, 1 mM EDTA, 15% formamide supplemented with PMSF, SUPERaseIn, and protease inhibitor as described above for lysis buffer) and 2.5 μL of the 100 μM ChIRP probe pool were added. In our experiments, the total volume of pre-cleaned sonicated lysate and ChIRP hybridization buffer was 12 ml per sample; for efficient hybridization, we distributed each sample in 6 ml aliquots and rotated in the hybridization over for 16 h at 37°C.

For enrichment of hybridized Zika virus RNA and associated proteins, per 1 mL of the initial sample, 250 μL of washed (washed 3 times in un-supplemented lysis buffer) Dynabeads MyOne Streptavidin C1 was added in each sample (in our study, 1 ml of beads was added per initial 4 ml lysis buffer sample), and rotated in the hybridization oven for 45 min at 37°C. Two 15 ml tubes representing one sample were placed on the magnetic stand, supernatants were removed, and beads were combined in 1 ml of ChIRP wash buffer (2x SSC, #AAJ60561AP, Fisher, MA, USA; 0.5% SDS supplemented with 1 mM PMSF just before use) in 1.5 ml low-binding Eppendorf tube. Afterward, beads were washed 5 times in 1 ml of the ChIRP wash buffer by rotating at 37°C for 5 min in the hybridization oven, placing for 4 min on the magnetic stand, and replacing the wash buffer. During the final wash, 10 μL of beads in the wash buffer were collected for RNA extraction, Zika-specific RT-qPCR and NGS to analyze the Zika RNA loads, entire virus genome coverage and NGS depth (see below and results).

For elution, beads were collected on the magnetic stand, resuspended in 600 μL of the ChIRP biotin elution buffer (12.5 mM biotin # B20656; 7.5 mM HEPES pH 7.9, #AAJ61275AK; 75 mM NaCl, # NC1071140; 1.5 mM EDTA, 0.15% SDS, 0.075% sarkosyl, # NC1386335; and 0.02% Sodium deoxycholate, #AAB2075914; all reagent are from Fisher, MA, USA), rotated at 25°C for 20 min, and at 65°C for 15 min. Beads were placed on the magnetic stand and eluents were transferred to a fresh tube, and beads were eluted again with 600 μL of the ChIRP biotin elution buffer. The two eluents were pooled (final 1200 μL) and residual beads were removed again using the magnetic stand. Afterward, 300 μL of trichloroacetic acid (#T0699; Millipore Sigma, MA, USA) (25% of the final elute volume) was added to the clean eluent, vortexed, and placed at 4°C overnight for precipitation. The next day, proteins were pelleted at 21,000 g at 4°C for 45 min. Supernatants were carefully removed, and protein pellets were washed once with ice-cold acetone (#A18-1; Fisher, MA, USA). Samples were spun at 21,000 g at 4°C for 5 min.

Acetone was removed, tubes briefly centrifuged again and after removal of residual acetone tubes were left to air-dry for 1 minute. Then, 60 µL of preheated 1× ChIRP lysis buffer containing 50 mM TEAB (# PI90114; Fisher, MA, USA) and 5% SDS (# BP2436-200; Fisher, MA, USA) were added to the protein pellet and gently mixed several times with 200 µL pipette tips. Samples were immediately put on dry ice and stored in –80°C until sent to the proteomics facility.

To validate the same viral RNA loads in all samples/experimental conditions at the time of formaldehyde RNA-protein fixation and ChIRP, RNA extracted during cell lysis (**input**) and elution steps (**enriched**) (see above) were tested with Zika virus-specific RT-qPCR assay and NGS to analyze the entire Zika virus genome coverage and NGS depth (**Supplemental Dataset 1**). The Zika virus-specific RT-qPCR assay was conducted as described above. For NGS, samples were brought to the final volume of 50 μL with 1x PBS and 5 μL Proteinase K (#FEREO0491; Fisher, MA, USA) and incubated at 55°C for 30 minutes. Then, RNA was extracted with the RNA Clean & Concentrator-5 Research Kit (ZYMO) and treated with the Invitrogen TURBO DNA-free Kit (#AM1907; Fisher, MA, USA). Afterward, 7 μL of RNA was used to produce cDNA with the Maxima H Minus First Strand cDNA Synthesis Kit (#FERK1652; Fisher, MA, USA) and random hexamers, following the manufacturer’s instructions. The single stranded cDNA (20 μL) was further used to prepare double-stranded DNA (dsDNA) using the NEBNext® Second Strand Synthesis (dNTP-free) Reaction Buffer (#B6117S; NEB, MA, USA) mixed with the DNA Polymerase I (#M0209S; NEB, MA, USA), E. coli DNA Ligase (#M0205S; NEB, MA, USA), RNase H (#M0297S; NEB, MA, USA), random hexamers (#N8080127; Fisher, MA, USA), and dNTPs mix (#18427088; Fisher, MA, USA). The dsDNA was used for library construction with the NEBNext® Ultra II DNA Library Prep Kit for Illumina (#E7645L; NEB, MA, USA) and NEBNext® Multiplex Oligos for Illumina® (96 Index Primers) (#E6609S; NEB, MA, USA) as we previously described [7,18,19]. Briefly, the libraries were constructed according to the manufacturer’s instructions, with 1:10 adaptor dilution (volume of adaptor:total volume), and following cleanup step of adaptor-ligated DNA without size selection. Individual libraries were quantified using Qubit dsDNA HS Assay Kit (#Q32856; Fisher, MA, USA) and Qubit 4.0 Fluorometer (Fisher, MA, USA). The peak size of the libraries was confirmed with the Agilent DNA 1,000 kit on Agilent 2100 Bioanalyzer (Agilent, CA, USA). Afterward, 20 ng of each barcoded library were pooled, quantified (Qubit), quality-checked (Agilent Bioanalyzer), and converted to moles: Molecular weight [nM] = Library concentration [ng/µL] / (Average library size x 650)/1,000,000). The pooled library was diluted to 2 nM in 10 mM TE buffer, denatured with 0.1 N NaOH, diluted to 14 pM, and paired-end 300 nt reads were generated with the MiSeq Reagent kit v3 600 cycle output (Illumina, CA, USA) on the MiSeq System (Illumina, CA, USA). To compare Zika virus RNA coverage and NGS depth, FASTQ files representing different samples/experimental conditions were trimmed for adaptor sequences and filtered for low-quality reads using Trimmomatic. Next, the trimmed FASTQ paired files were aligned to the Zika virus reference sequence H/PF/2013 [GenBank: KJ776791.2] using Burrows-Wheeler Aligner (BWA) [20] and output BAM files were used to quantify and visualize the virus genome coverage and NGS depth with the in-house R script (https://github.com/itrus/bash-scripts-NGS/tree/main) [7,18,19].

Preparation and mass spectrometry analysis of ChIRP samples were done in The Proteomics Facility, Network Biology Collaborative Centre, Lunenfeld-Tanenbaum Research Institute, Toronto. For the S-Trap (ProtiFi, NY, USA) [21,22] the whole volume containing ChIRP-precipitated proteins were used. The material was reduced with 20 mM DTT for 10 minutes at +95°C and alkylated with 40 mM iodoacetamide for 30 minutes in the dark. Phosphoric acid was added to achieve a final concentration of 1.2%. To each 27.5 µl of acidified lysate, 165 µl of S-Trap protein binding buffer (90% methanol, 100 mM TEAB) was added. The resulting mixture was passed through the micro column at 4,000 × g, and the column was washed four times with the S-Trap protein binding buffer. Each sample was digested with 1 µg of trypsin (in 20 µl of 50 mM TEAB) for 1 hour at +47°C. Prior to elution, 40 µl of 50 mM TEAB (pH 8) was added to the column. Peptides were eluted by centrifugation at 4,000 × g. Peptides were eluted two more times with 40 µl of 2% formic acid and 40 µl of 50% acetonitrile + 2% formic acid. Eluted peptides were dried down and stored at –40°C.

For data-dependent acquisition (DDA) LC-MS/MS, digested peptides were analyzed using a nano-HPLC coupled to MS. Half of the sample was used for DDA. Coated Nano-spray emitters were generated from fused silica capillary tubing, with 75µm internal diameter, 365µm outer diameter and 5-8µm tip opening, using a laser puller (Sutter Instrument Co., model P-2000, with parameters set as heat: 280, FIL = 0, VEL = 18, DEL = 2000). Nano-spray emitters were packed with C18 reversed-phase material (Reprosil-Pur 120 C18-AQ, 1.9 µm) resuspended in methanol using a pressure injection cell. The sample in 5% formic acid was directly loaded at 400 nl/min onto a 75 µm x 15 cm nano-spray emitter. Peptides were eluted from the column with an acetonitrile gradient generated by an Vanquish Neo UHPLC System and analyzed on an Orbitrap Fusion Lumos Tribrid. The gradient was delivered at 200 nl/min from 3.2% acetonitrile with 0.1% formic acid to 35.2% acetonitrile with 0.1% formic acid using a linear gradient of 90 minutes. This was followed by a 3 min gradient from 35.2% acetonitrile with 0.1% formic acid to 79.2% acetonitrile with 0.1% formic acid. Afterward, there was a 5.8 min wash with 79.2% acetonitrile with 0.1% formic acid, and the column was equilibrated at 400nl/min for 6.4min. The total DDA protocol was 99 minutes. The MS1 scan had an accumulation time of 50 ms within a mass range of 400–1500Da, using orbitrap resolution of 120000, 60% RF lens, AGC target of 125% and 2400 volts. This was followed by MS/MS scans with a total cycle time of 3 seconds. Accumulation time of 50 ms and 33% HCD collision energy was used for each MS/MS scan. Each candidate ion was required to have a charge state from 2-7 and an AGC target of 400%, isolated using an orbitrap resolution of 15,000. Previously analyzed candidate ions were dynamically excluded for 9 seconds.

Mass spectrometry RAW data files (**Supplemental Dataset 2**) were converted using ThermoRawFileParser GUI –V 1.7.1 (https://github.com/compomics/ThermoRawFileParserGUI) to the mzML format compatible with FragPipe. The RAW files were analyzed using FragPipe v18.0 (**Supplemental Dataset 2**), which utilized MSFragger v3.5, Philosopher v4.4.0, and IonQuant for peptide searching in the raw files [23–25]. The LFQ-MBR workflow was used, with the *Chlorocebus sabaeus* Proteome ID UP000029965 (https://www.uniprot.org/proteomes/UP000029965) database with appended Zika virus H/PF/2013 [GenBank: KJ776791.2], decoys, and contaminants sequences. Data from the output “combined_protein.tsv” file (**Table S1B; Supplemental Dataset 2**) were used for interactome analysis described below.

### Identification of Extended, ZAP-independent, and ZAP-dependent Zika virus RNA interactomes from ChIRP-MS results

The Zika virus RNA-interacting proteins were scored with the Mass spectrometry interaction STatistics (MiST) analysis which was previously applied for ChIRP-MS [17,26,27]. We used MiST default parameters; for specificity, we also selected the Singleton Filtering MiST option to exclude proteins with spectral counts in only one biological replicate and quantify proteins with MiST score 0.75 or above [17,26,27]. We also excluded all proteins which had at least a single spectral count in at least one of biological replicate in mock-infected wild-type or ZAP-KO cells [17,28].

We applied three analytical strategies to examine ChIRP-MS data. For the first analytical strategy, using the raw ChIRP-MS data with total 2,269 proteins (**Table S1B)**, we identified proteins specifically associated with Zika virus RNA in VERO-ZAP-WT cells. We defined the resultant list of enriched proteins as “extended” Zika virus RNA interactome in VERO-ZAP-WT cells. For the second analytical strategy, we identified proteins specifically interacting with Zika virus RNA in VERO-ZAP-KO cells. We defined the resultant list of enriched proteins as “ZAP-independent” Zika virus RNA interactome. We defined this interactome as ZAP-independent because associated cellular proteins were discovered in VERO-ZAP-KO cells which shows ZAP-independent nature of interactions. Finally, for the discovery of “ZAP-dependent” Zika RNA interactome in VERO-ZAP-WT cells, in addition to the above specificity exclusion criteria in MiST settings and in mock-infected cells, we excluded proteins which had at least one spectral count in at least one of the biological replicates in VERO-ZAP-KO cells infected with Zika virus.

For ZAP-independent and ZAP-dependent Zika virus RNA interactomes, we performed GO enrichment analyses of the identified proteins using The Gene Ontology Resource (https://geneontology.org/) which connects to the analysis tool from the PANTHER Classification System [29–31]. For analysis, GO aspect molecular function was selected. As recommended, a custom reference list containing all proteins’ IDs identified in the specific mass spectrometry run (**Tables S1B**) was added after initial analysis on The Gene Ontology Resource and the analysis was re-run on the PANTHER website. The significance *P* values of GO terms were calculated by Fisher’s exact test and adjusted by FDR.

### Statistics

We used GraphPad PRISM 9 software. For the comparison of the number of virus-positive cell counts, and viral RNA loads between wild-type and ZAP-KO cells we used an unpaired t-test. For the comparison of Western blot protein expression, we used one-way ANOVA multiple comparisons. *P*-value < 0.05 was considered statistically significant.

